# Targeted disruption of *Pparγ1* promotes trophoblast endoreplication in the murine placenta

**DOI:** 10.1101/2020.05.28.120691

**Authors:** Takanari Nakano, Hidekazu Aochi, Masataka Hirasaki, Yasuhiro Takenaka, Koji Fujita, Hiroaki Soma, Hajime Kamezawa, Takahiro Koizumi, Akihiko Okuda, Takayuki Murakoshi, Akira Shimada, Ikuo Inoue

## Abstract

In murine placentas, peroxisome proliferator-activated receptor (PPAR) γ1, a nuclear receptor, is abundant at the late stage of pregnancy (E15–E16), but its functional roles are still elusive because PPARγ-full knockout embryos die early (E10). We generated mice disrupted in only *Pparγ1*, one of the two major mRNA splicing variants of PPARγ1. *Pparγ1-* knockout embryos developed normally until 15.5 dpc, but their growth was retarded thereafter and they did not survive. At 15.5 dpc, in the wild-type placentas, intense PPARγ-immunostaining was detected in sinusoidal trophoblast giant cells (sTGCs), a cell lineage that coordinates the maternal blood microcirculation in the labyrinth, whereas they were absent in the knockouts. Although *Pparγ1*-knockout placentas were normal in morphology, we observed severely dilated maternal blood sinuses in the labyrinth. The *Pparγ1*-knockout sTGCs had abnormally large nuclei, an enhanced endocycling phenotype, indicating insufficient differentiation. RNA-sequencing of the placentas showed increased expression of genes coding for nucleosome assembly factors. Labyrinthine gene expressions for atypical E2Fs and cyclin E, key drivers for endocycling, were increased >3-fold. These findings suggested that PPARγ1 plays a key role in endocycle termination.

## Introduction

Peroxisome proliferator-activated receptor (PPAR) γ, a nuclear receptor, plays key roles in a number of cellular processes such as cell differentiation (Lefterova et al., 2014) and the catabolism of nutrients (Grygiel-Górniak, 2014). PPARγ is mainly comprised of two isoforms, PPARγ1 and PPARγ2. PPARγ2 is predominantly present in adipose tissues (Poulsen et al., 2012), whereas PPARγ1 is expressed ubiquitously. The placenta highly expresses PPARγ1 (Takenaka et al., 2013), which is involved in embryonic development although the mechanisms involved are currently unknown.

PPARγ is required to carry fetuses to term (Barak et al., 1999). PPARγ-knockout (KO) embryos died before embryonic day (E)10.5 with a widely disrupted feto–maternal vascular network in the labyrinth of the chorio-allantoic placenta (Barak et al., 1999; Kubota et al., 1999). Although PPARγ is abundant in the placenta in the late stage of pregnancy (Barak et al., 1999; Nadra et al., 2010), the critical cell lineages, cellular processes, and molecular pathways regulated by PPARγ1 remain elusive (Barak et al., 2008; Segond et al., 2013; Akshaya et al., 2015).

The labyrinth of the placenta provides a large surface area for feto–maternal gas and nutrient exchange in rodents, where the fetal and maternal vascular structures are closely associated with each other (Adamson et al., 2002). Labyrinthine trophoblast giant cells (TGCs) secrete a wide array of hormones and paracrine factors (Wynne et al., 2006; Marano and Ben-Jonathan, 2014) to ensure correct vasculature development in the placenta. At least four different subtypes of TGCs are present within the mature placenta (Hu and Cross, 2010). Among them, sinusoidal TGCs line up at the surface of the maternal capillaries for development and maintenance (El-Hashash et al., 2010).

TGCs undergo chromosomal endoreplication, a phenomenon where the genome replicates without entering mitosis to become highly polypoid cells (>1000C) (Fox and Duronio, 2013). Deprivation of fibroblast growth factor 4 initiates the anomalous cell cycle (Ullah et al., 2008). So far, several key cascade molecules, such as cyclin E and atypical E2Fs (E2F7/8) have been revealed (Geng et al., 2003; Chen et al., 2012; Zielke et al., 2013), but it is unknown how the cells regulate endocycle termination.

In expectation of obtaining a milder phenotype of PPARγ1 disruption, we generated a mouse line lacking expression of *Pparγ1*, one of the two mRNA splicing variants coding for PPARγ1. *Pparγ1-KO* embryos developed normally until 15.5 days post-coitum (dpc), but revealed growth restriction thereafter and did not survive. In the present study, we examined the functional role(s) of *Pparγ1* in the late stage of pregnancy in transgenic mice to further understand the role of PPARγ1 in embryonic development.

## Results

### Generation of *Pparγ1- or Pparγ1-knockout* mice

*Pparγ1* and *Pparγ1sv* (for the exon structure, see Fig. S1) are splicing variants coding for murine PPARγ1. The FANTOM database (http://fantom.gsc.riken.jp/zenbu/) indicated that both isoforms are expressed in the placenta, with *Pparγ1* being predominant (Fig. S2). We confirmed the expression of both isoforms and higher expression of *Pparγ1* than that of *Pparγ1sv* using quantitative reverse transcriptase-PCR (RT-qPCR) with calibrated reference standards (Virtue et al., 2010; Takenaka et al., 2013) in C57BL/6J placentas. The expression levels had peaks at 16.5 dpc (Fig. 1). *Pparγ*-full deletion produced no available conceptuses later than E10.5 (Barak et al., 1999; Kubota et al., 1999). Because *Pparγ2* produced no detectable amount of transcripts in the placenta (Barak et al., 1999, also see Fig. 1A), the defect in *Pparγ*-full knockout fetuses was exclusively due to the loss of expression of PPARγ1. Accordingly, *Pparγ*-full knockout can be assumed to cause the absence of PPARγ1. Therefore, we generated mice that expressed either *Pparγ1* or *Pparγ1sv* isoforms in the hope for reducing the pathological severity, which might allow assessment of the physiological importance of PPARγ1 around the peak expression of the coding genes in the placenta (Fig. S3A).

**Figure 1.**
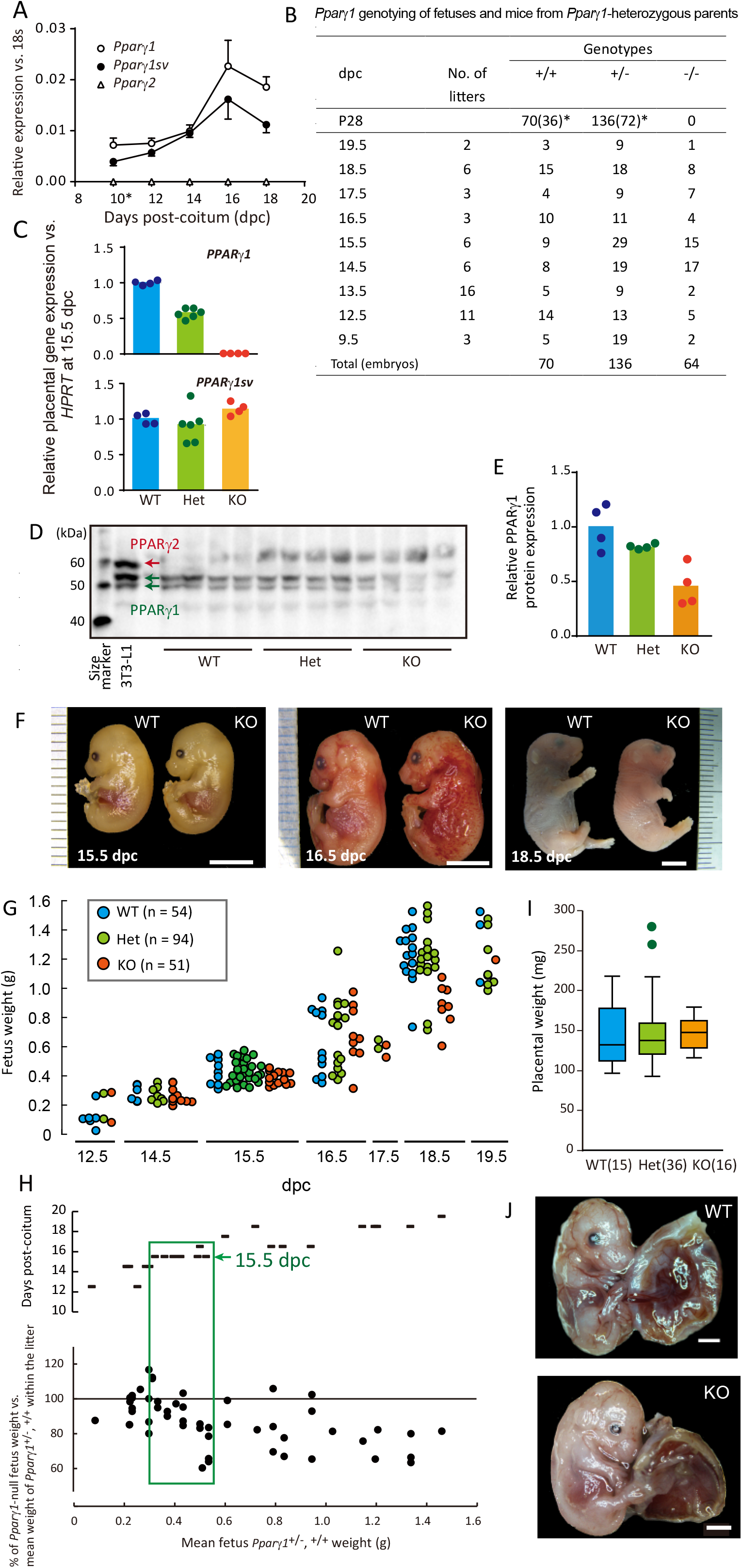
*A*, Semi-quantitative gene expression analysis of *Pparγ* variants in the placentas of C57BL/6J mice from 10 to 18 dpc. Ribosomal *18S* was used to normalize expression levels. Each plot represents the mean of n = 5 biological replications and SEM. *B*, Distribution of *Pparγ1* genotypes in fetuses and adult mice from the *Pparγ1*-heterozygous mating at various time points during pregnancies. *Pparγ1*-KO mice were present until the term, but no *Pparγ1*-KO pups were obtained. *Numbers of females in parentheses. *C-E*, Gene expression of *Pparγ1 (upper)* and *Pparγ1sv* (*bottom*) (*C*) and PPARγ abundance *(D* and *E*) in placentas at 15.5 dpc. *E* shows a comparison of the intensity between the genotypes (one-way ANOVA, *F*(2,9) = 11.78, p = 0.003). For each lane, 10 μg of placental protein extract was loaded. Bars indicate mean values. As a positive control, 6 μg protein of differentiated 3T3-L1 nuclear fraction was loaded. *F*, Representative gross appearances of WT (+/+) and *Pparγ1*-KO (-/-) fetuses at the indicated time points. Bars indicate 5 mm. *G*, Body weight of fetuses at the time points indicated. *H*, Y-axis *(upper)* indicates dpc of the female mice examined; the Y-axis *(bottom)*, percentage of *Parγ1*-KO fetus body weight normalized against the WT and heterozygotes at different time points. X-axis, the mean fetus body weight for the WT and heterozygotes. Each bar plot in the upper panel shows an individual pregnant mouse. These were aligned according to their the mean values. Each plot (*circles*) shows a normalized percentage of individual *Parγ1*-KO fetuses by weight (*bottom*). The green box indicates the period where growth restriction began. At 15.5 dpc, mean fetus weight with the WT and heterozygotes ranged from approximately 0.3 to 0.5 g. The growth restriction was obvious when the mean weight was more than 0.4 g. *I*, Placental weights observed at 14.5 or 15.5 dpcs. The numbers examined are shown in parentheses. *J*, Representative gross embryo appearances at 15.5 dpc. Het indicates *Pparγ1*^+/-^.

### *Pparγ1*-KO mice are embryonic lethal

Genotyping of pups by PCR (Fig. S3B) showed that no *Pparγ1*-KO mice were born live. *Pparγ1sv-KO* mice maturated and were fertile after backcrossing to B6, although a proportion of the *Pparγ1sv*-KO embryos died (Tables S1, S2). *Pparγ1sv*-KO mice had lower body weight at post-weaning and sometimes through life in males (Fig. S4). Accordingly, *Pparγ1*, rather than *Pparγ1sv*, was crucial for carrying the fetuses to term; thus, we examined the phenotypes of *Pparγ1*-KO conceptuses.

### Intrauterine growth restriction of *Pparγ1*-KO fetuses began at 15.5 dpc

The genotypes of the fetuses from *Pparγ1*^+/-^(heterozygote, Het)-mating yielded a Mendelian ratio of approximately 1:2:1 for wild-type (WT), Het, and KO, respectively (Fig. 1B). *Pparγ1* expression (Fig. 1C) and protein levels (Figs. 1D-E, for western blotting validation, see Fig. S5) were in accordance with the gene dosage at 15.5 dpc. From gross examinations, *Pparγ1*-KO fetuses appeared to grow normally until 15.5 dpc (Fig. 1F, *left*), but suffered massive hemorrhage in the skin at 16.5 dpc (Fig. 1F, *center*), and showed apparent smaller body size at 18.5 dpc (Fig. 1F, *right*). Measurement of fetus weight indicated that the intrauterine growth restriction of *Pparγ1-KO* fetuses started around 15.5 dpc (Figs. 1G-H), which was close to the time when *Pparγ1* expression reached its peak (Fig. 1A). The KO fetuses kept growing until term but at a reduced rate (about 20%). The heterozygotes grew normally and were fertile and lacked external symptoms of disease when mature (Figs. S6A-B).

### *Pparγ1*-KO placentas formed normally

Intrauterine growth restriction often originates from insufficient formation of the placenta (Lean et al., 2017). We weighed the placentas but found no significant difference between the genotypes (Fig. 1I). In addition, we did not observe any consistent morphological differences macroscopically (Fig. 1J).

### Abundant PPARγ1 expression in sinusoidal trophoblast giant cells

Developmental defects in *Pparγ*-full KO labyrinth indicated that PPARγ1 is expressed in labyrinthine trophoblast cells (Barak et al., 1999), but it was uncertain which lineages expressed the protein and were affected. To identify the cell types, we immunostained sections obtained from placentas at 15.5 dpc for PPARγ and found that large, round immunostaining puncta were observed in the labyrinthine region of WT placentas (Figs. 2A-I). The large nuclei were confirmed by methyl green counterstaining (Fig. S7A), showing that ‘sinusoidal’ TGCs were positive for PPARγ1 (El-Hashash et al., 2010; Hu and Cross, 2010).

**Figure 2.**
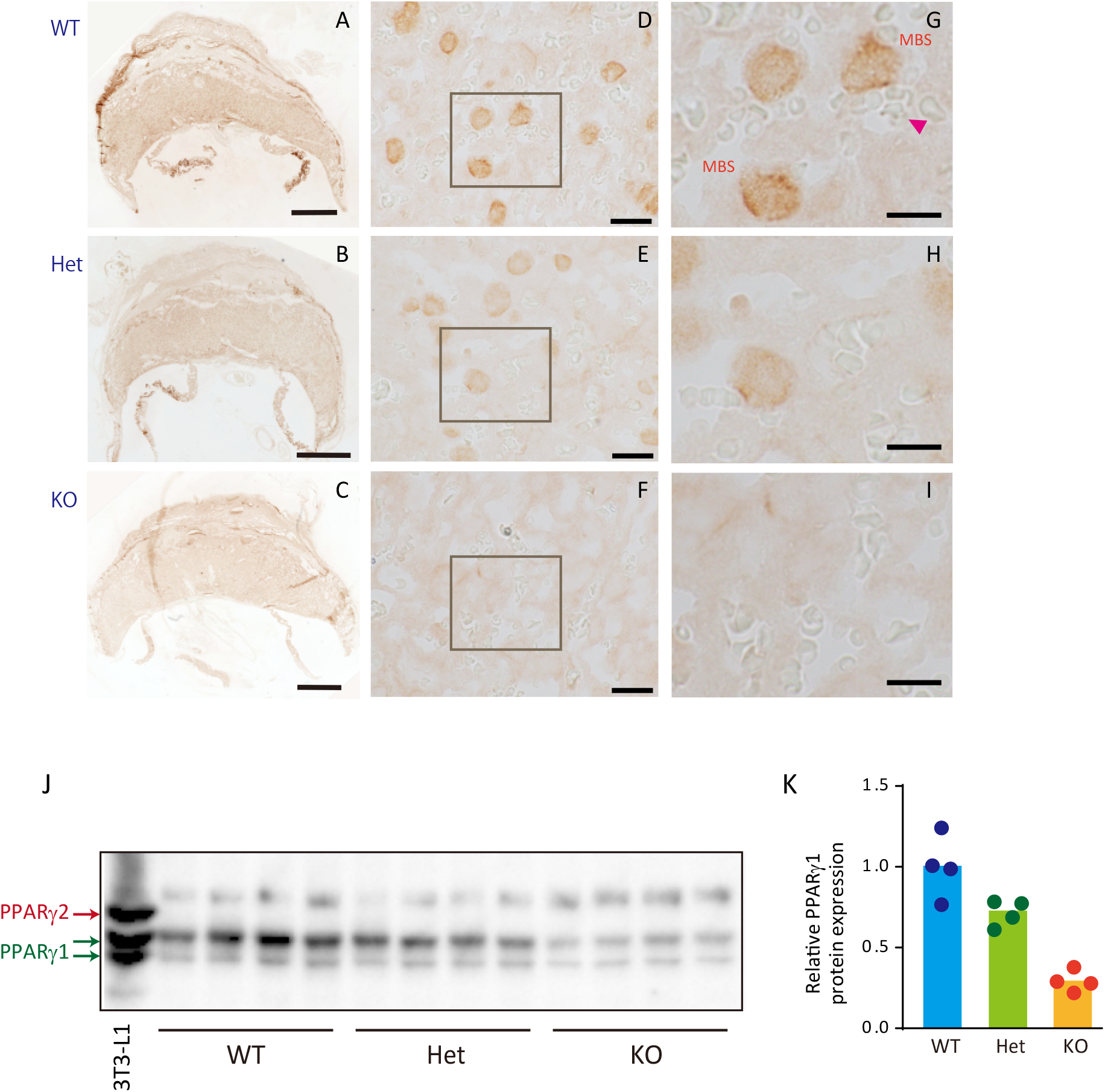
*A–I*, PPARγ expression in the labyrinth. *A–C*, PPARγ-immunostained placental sections. Bars = 1 mm. *D-F*, labyrinth; bars = 20 μm. *G–I*, cropped images; bars = 10 μm. *J* and *K*, MBS, maternal blood sinus; arrow head, red blood cells. Labyrinthine PPARγ protein abundance determined using Western blotting (*J*) and quantified according to the intensity (*K*). WT, Het, and KO indicate the *Pparγ1* genotypes.

The immunostaining intensity was approximately half of that in heterozygotes (Fig. S7B), and disappeared in *Pparγ1-*KO. Western blotting showed that PPARγ1 abundance was approximately one-third in the labyrinth of *Pparγ1*-KO (Figs. 2J-K), and the residue may have originated from *Pparγ1sv*. Apparent PPARγ1 expression was not observed in other TGC species (Fig. S8) (For TGC lineages, see Hu and Cross, 2010).

We found intense PPARγ-staining in the cytosol of endodermal cells of the yolk sac membrane (Figs. S8I-K). PPARγ1 was almost absent in *Pparγ1-*KO, indicating that PPARγ1 in the endodermal cells originated from *Pparγ1*. Barak et al. found pale yolk sacs at E10–11.5 of *Pparγ*-full KO conceptuses, suggesting involvement in the physiology or development of the yolk sac in the present study (Barak et al., 1999), although *Pparγ1*-KO did not affect structural development appreciably (Figs. S8I-J).

### Larger nuclei of sinusoidal TGCs in the labyrinth of *Pparγ1* KO placentas

We then further examined *Pparγ1-KO* placentas histologically. Periodic acid-Schiff (PAS) staining showed that all the major characteristic layers were present in *Pparγ1-mutant* placentas at 15.5 dpc (Figs. 3A-C) with normal labyrinthine maturation in size [% labyrinth vs. labyrinth + junctional zone in area; one-way ANOVA *F*(2,6) = 2.003, *p* = 0.25]. Hematoxylin and eosin (H&E) staining of the placentas revealed that sinusoidal TGCs had abnormally larger nuclei in *Pparγ1*-KO placentas with inappropriate location and with occasionally aggregation (Figs. 3G–L). Examination of nuclear morphology showed that the frequency of binucleation in sinusoidal TGCs was not different appreciably, indicating that the enlarged nuclei were not due to repeated endomitosis. Feulgen staining (Figs. 3N, O) and the following semi-quantification of chromosome abundance showed that normalized chromosome density was almost equal between the WT and KO (Fig. 3P). Accordingly, chromosome content calculated from the data in Figures 3M and 3P was estimated to be about 3-fold greater for the chromosome abundance in *Pparγ1*-KO (Fig. 3Q). On the other hand, there was no significant difference in the number of sinusoidal TGCs (Fig. 3R).

**Figure 3.**
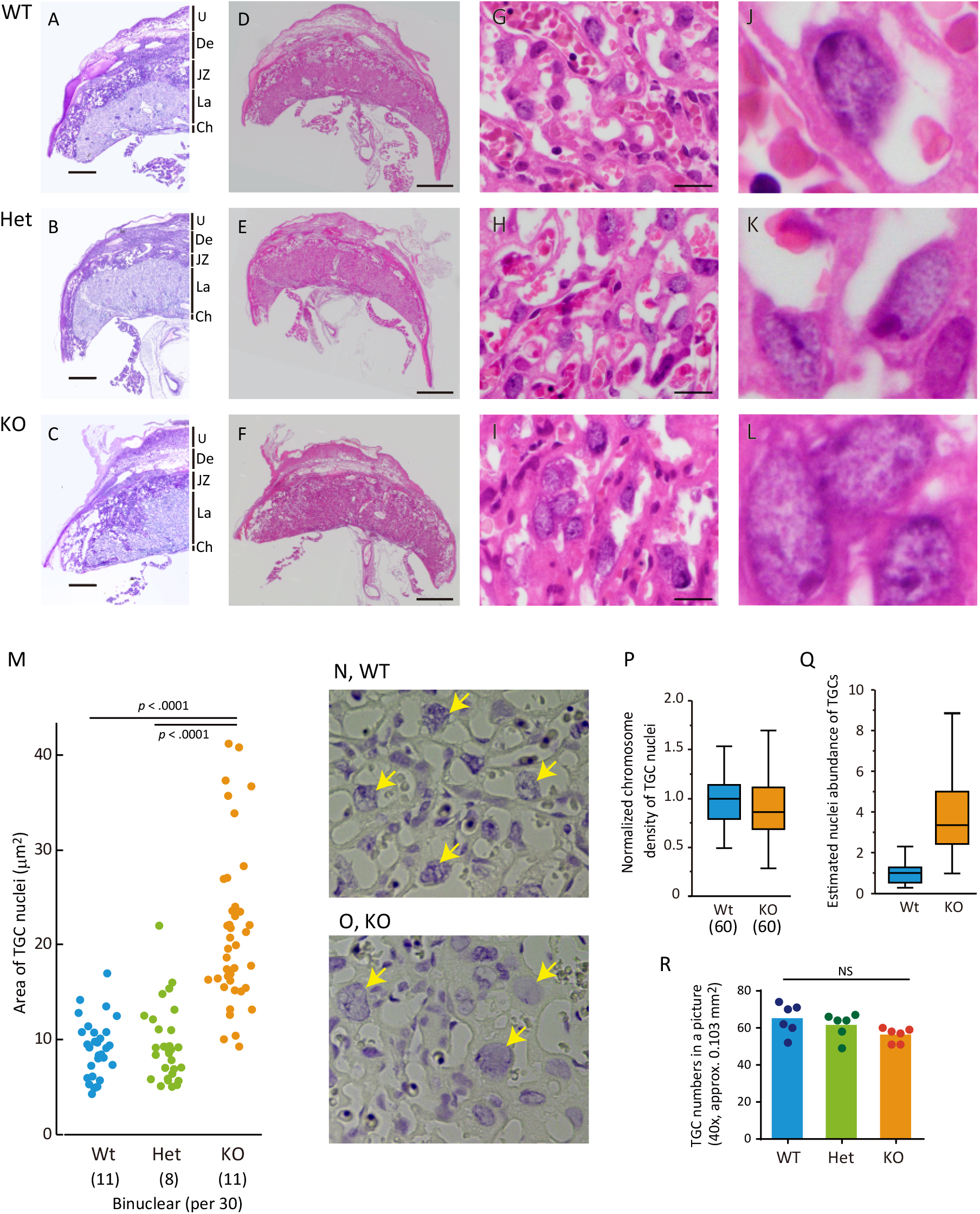
Histologic examination of placentas at 15.5 dpc. Upper panels (*A, D, G*, and *J*), *Pparγ1*-WT; middle panels (*B, E, H*, and *K*), heterozygotes; lower panels (*C, F, I*, and *L*), KO. *A–C*, Low magnification view of PAS and hematoxylin-stained sections. *D–F*, Sections stained with H&E. Panels *A–F* show that placentas with the *Pparγ1*-KO allele developed normally in terms of scale and overall structure. U, uterus; De, decidua; JZ, junctional zone; La, labyrinth; Ch, chorion. *G–I*, Labyrinth and respective cropped images (*J–L*). In the KO-labyrinth, a few sinusoidal TGCs gathered occasionally, but the vessels were disrupted. *AC*, bars = 0.5 mm; *D-F*, bars = 1 mm; *G-I*, bars = 20 μm. *M*, Difference in the area of sinusoidal TGC nuclei (n = 30). The parentheses in the bottom indicate the number of cells with bi-nuclear structures. *P* value was obtained with the Steel–Dwass test. *N* and *O*, Feulgen staining of the labyrinth. *P*, Relative chromosome density of TGCs. *Q*, Estimated chromosome abundance. Assuming that chromosome density was the same between the genotypes (*P*), the chromosome abundance is proportional to the size. Data used in *M* was utilized for the volume estimation. *R*, Numbers of TGCs in the labyrinth was not different between the genotypes.

### Disturbed labyrinthine architecture in *Pparγ1*-KO placenta

In addition to the enlarged sinusoidal TGC nuclei, H&E staining showed frequently dilated, ruptured blood spaces, especially in the maternal blood sinuses. We measured the blood spaces and found dilation especially in the maternal blood sinuses (Figs. 3D-F).

Monocarboxylate transporter-1 (MCT1) is a marker for type-I syncytiotrophoblasts that face the maternal sinuses comprising feto–maternal interfaces (Moreau et al., 2014). To further reveal the structural alterations with *Pparγ1-*KO, we stained for MCT1, showing that each sinusoidal TGC was surrounded by type-I syncytiotrophoblasts in WT (Fig. 4A). On the other hand, the structure was compromised in the mutants, especially in *Pparγ1*-KO (Figs. 4B-C).

**Figure 4.**
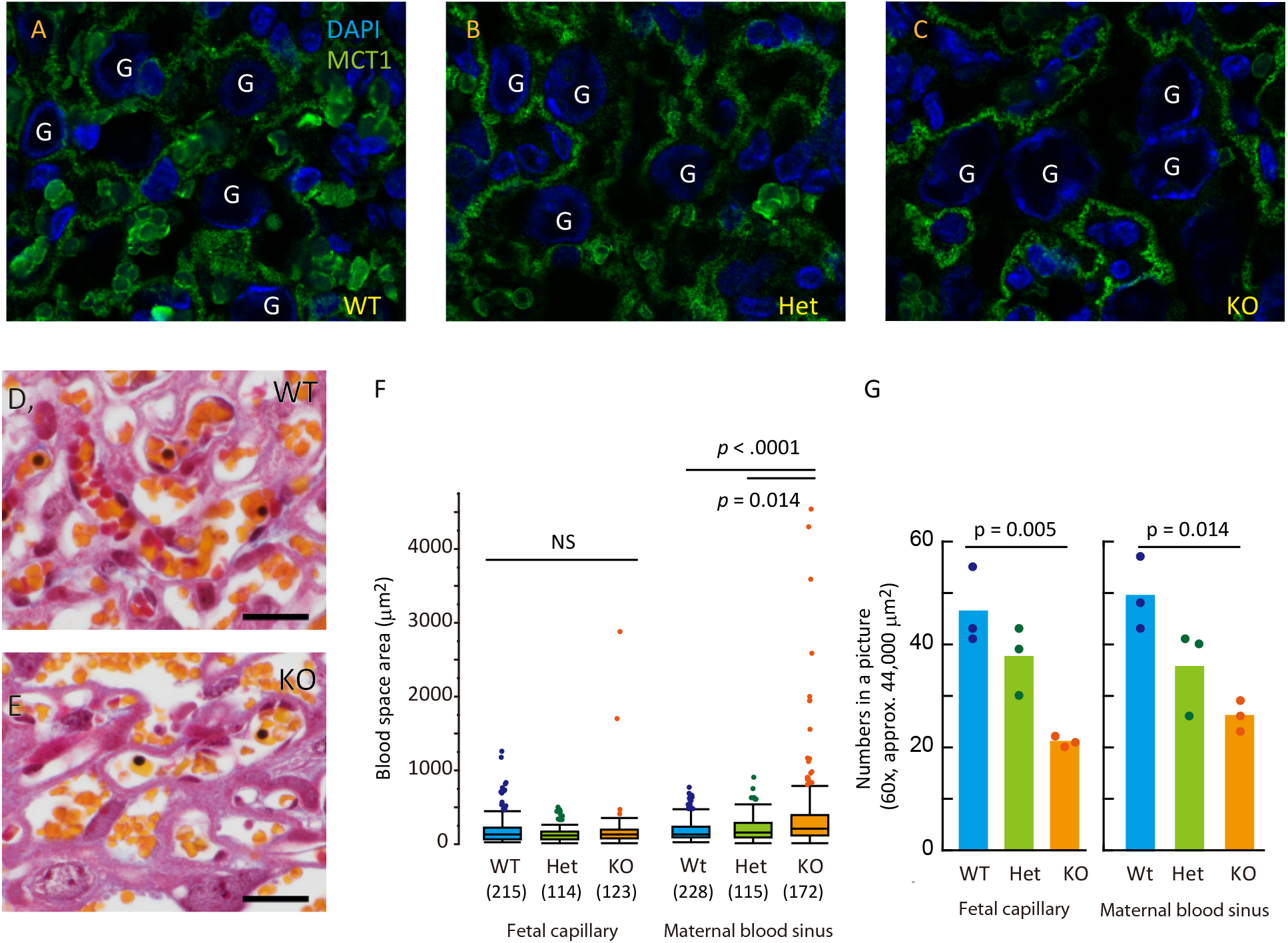
*A–C*, Immunohistochemical detection of MCT-1, a marker for syncytiotrophoblast I. Genotypes are indicated. G, nuclei of TGCs. *D* and *E*, Masson trichrome staining for the labyrinth. Fewer blood cells were observed in the KO. bars = 20 μm. *F* and *G*, Comparison of blood vessel area and vessel number, respectively, in the labyrinth between genotypes. Three sections were stained for Masson trichrome and images obtained with a 40× objective (*e.g., D and E*). *F*, Area of vessels in the labyrinth was measured randomly. The numbers in parentheses indicate the number of vessels examined. Differences between the genotypes was analyzed using Steel–Dwass test (*F*). *G*, Bars indicate mean sinusoidal TGC numbers in an image (60× objective). A dot shows the average of three images from a section. n = 3 biological replications. one-way ANOVA, *F*(2,6) = 14.46, *p* = 0.005 for *left, F*(2,6) = 9.54, *p* = 0.014 for *right* in *G*. WT, Het, and KO indicate *Pparγ1* genotypes.

We measured the blood space area of sections stained with Masson trichrome (Figs. 4D-E) and determined that the area of the maternal blood sinuses, but not the fetal capillaries, was occasionally much larger in the mutants, indicating aberrant and decreased vascularization of the maternal blood spaces (Fig. 4F). In addition, the number of the blood vessels was lower with the mutation (Fig. 4G). Such structural impairments were also confirmed by scanning electron microscopy (Fig. S9).

The severity was probably dependent on PPARγ1 abundance. Thus, we examined *Pparγ1sv-KO* placentas histologically and compared them with *Pparγ1*-KO placentas. Similar but slightly dilated blood capillaries were observed in *Pparγ1sv*-KO labyrinths (Fig. S10). In addition, an apparent increase in size was observed with *Pparγ1sv*-KO, and the nuclei in *Pparγ1sv*-KO had sparse hematoxylin staining in the inner regions similar to *Pparγ1*-KO (Fig. 3). The localization was appropriate, facing the maternal blood sinuses without aggregation with sinusoidal TGCs. Accordingly, these findings suggested that the severity was in accordance with the total PPARγ1 abundance in sinusoidal TGCs.

### Insufficient support for maternal sinus development with *Pparγ1*-KO sinusoidal TGCs

To look into histological changes, especially in sinusoidal TGC localization and the structure of the feto–maternal interfaces in the labyrinth, we performed transmission electron microscopy of the labyrinth area. In the WT, sinusoidal TGCs faced the maternal blood (Fig. 5A). Rich feto–maternal interfaces were obvious with crowded blood spaces (Figs. 5A, B, S11A). For example, a sinusoidal TGC was located at the junction of the blood spaces with a microvillus structure on the surface (Figs. 5C, D). In contrast, KO sinusoidal TGCs did not face the maternal blood sinuses but occasionally adjoined (Figs. 5E, S11B). Furthermore, erythrophagocytosis was found through the KO-labyrinth, presumably via KO-sinusoidal TGCs (Fig. 5F).

**Figure 5.**
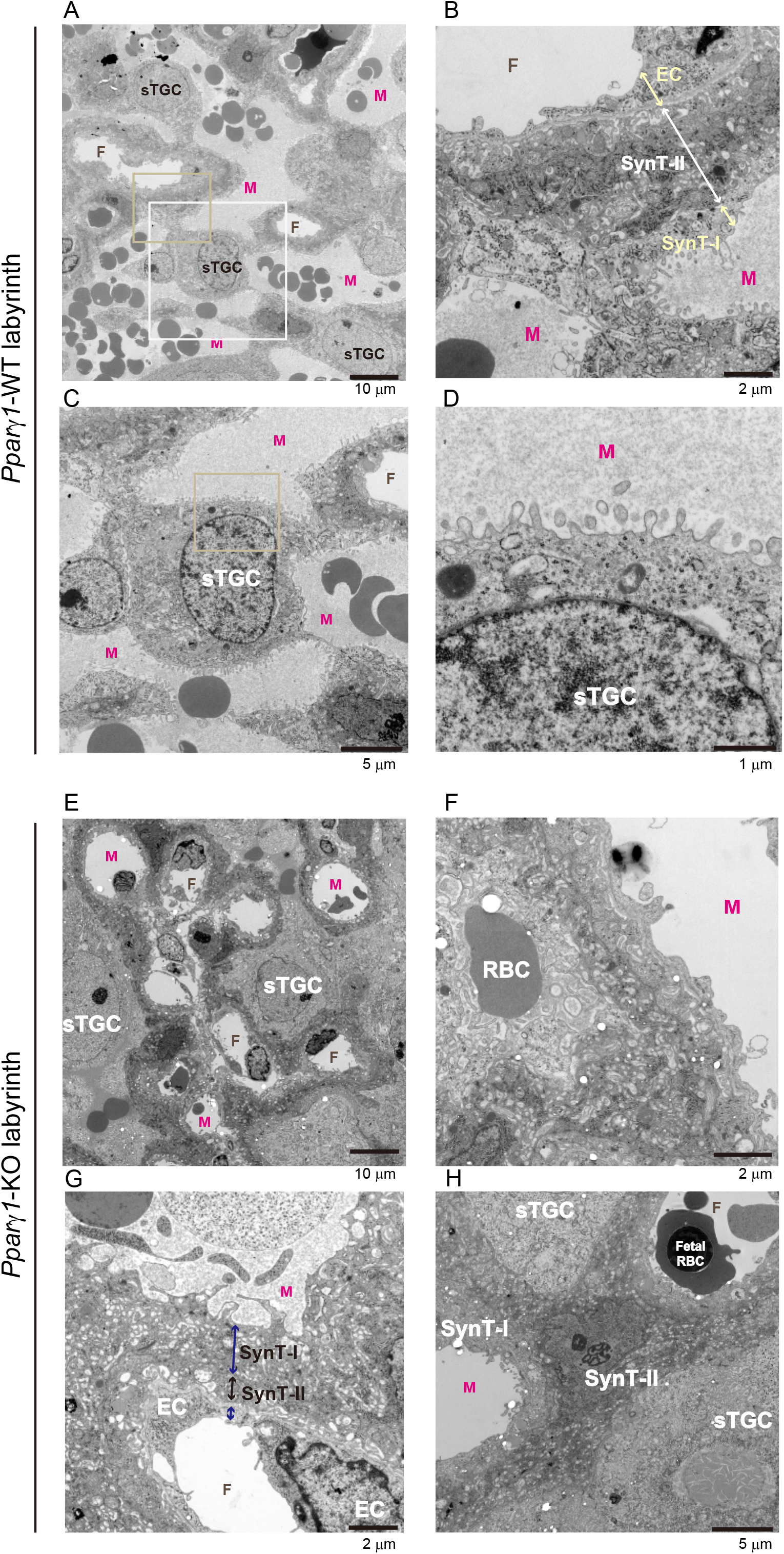
*A–D*, WT; *E–H*, KO. A shows the spatial localization of sinusoidal TGCs (sTGCs) and blood spaces in the section. Developed and rich feto–maternal boundaries are observed. *B*, a Representative feto–maternal boundary cropped from panel *A* (*yellow rectangle*). Arrows indicate each cell layer for the boundary. *C*, Shows a sinusoidal TGC that is located at the junction of the blood spaces cropped from panel *A* (*white rectangle*). *D*, Microvillus of the sinusoidal TGCs in *C* (cropped area is shown as a square in *C*). *E* The spatial localization of sTGCs with dilated blood spaces in the section. *F*, A red blood cell (RBC) was phagocytosed. *G*, Feto–maternal boundary found in a KO-labyrinth section. Such a boundary was rarely observed. Arrows indicate each cell layer for the boundary. *H*, A type-II syncytiotrophoblast (synT-II) impeded a feto–maternal interface by staying at the boundary. TGC, trophoblast giant cells; M, maternal blood sinus; SynT-I, type-I syncytiotrophoblast; EC, endothelial cell; F, fetal blood capillary. Bars in *A* and *B*, 10 μm; 2 μm in *C* and *D*.

When we compared feto–maternal boundaries in the labyrinth and observed the characteristic three cell layers in the WT (Figs. 5B, S11C); in the *Pparγ1*-KO, such a characteristic thin layer was rarely found (Figs. 5E, S11B). On the other hand, in addition to the sinusoidal TGCs, we found compromised type-II syncytiotrophoblasts (Figs. 5G-H) that were composed of the center of the three characteristic layers (Figs. 5E, S11B). In Figure 5G, the type-II syncytiotrophoblast layer was thin: In Figure 5H, one of the cells did not form a thin layer for the boundaries.

We also found that the type-I syncytiotrophoblasts, which face the maternal blood sinuses and contribute to nutrient uptake from the maternal blood (Fig. S11C), had poor villus development in the KO (Fig. S11D). These findings suggested that abnormality of sinusoidal TGCs affected the functionality of neighboring cells (Natale et al., 2009).

### RNA-sequencing indicated increased nucleosome assembly in *Pparγ1*-KO placentas

Sinusoidal TGCs undergo endoreplication, which is pivotal to placental and fetal development (Geng et al., 2003; Parisi et al., 2003). Chen et al. showed that promoted endoreplication resulted in abnormally enlarged nuclei in TGCs (Chen et al., 2012). To determine whether enlarged nuclei in *Pparγ1*-KO are related to endoreplication progression as reported, we performed RNA-sequencing analysis of the placentas. The analysis again confirmed the absence of *Pparγ1* expression in exon A1-deleted mice (Figs. S12A-B). There were 1126 (WT > KO) and 319 (KO > WT) genes that were significantly up-regulated and down-regulated, respectively (more than 1.3-fold difference, *p* < 0.05; n = 4 for each group), in the placentas (Fig. 6A). Gene ontology analysis indicated that the genes with increased expression levels were associated with DNA replication, nucleosome assembly, and transcription inhibition (Fig. 6B; for the table, see Fig. S12C). We further examined the individual genes in these groups and found higher expression levels among histone-related genes in the *Pparγ1-KO* (Fig. S12D), indicating that chromosome replication activity was maintained at a high level in the KO.

**Figure 6.**
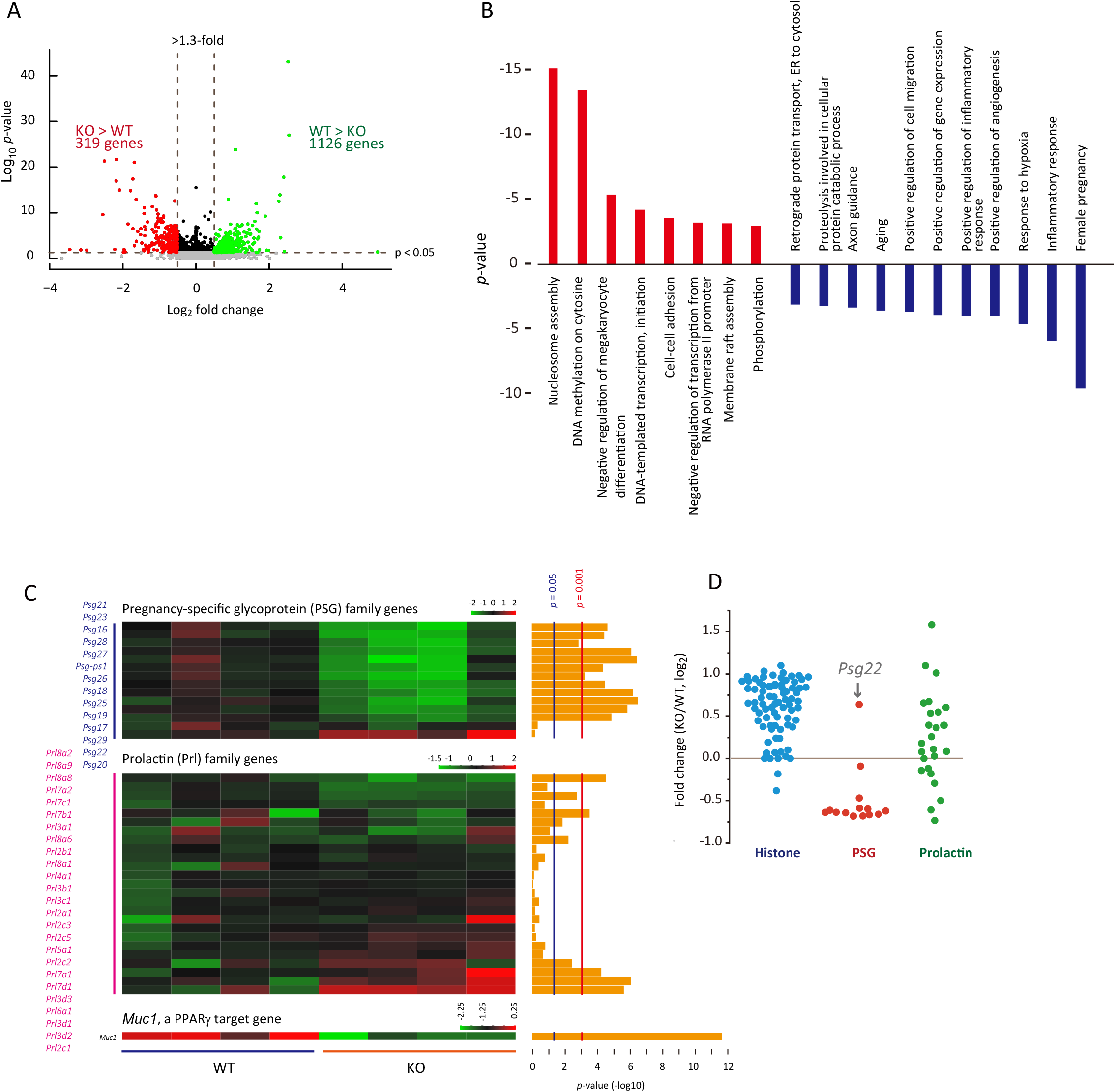
RNA-sequencing analysis of placentas. *A*, Volcano plot for the RNA-sequencing analysis. Gray, *p* ≥ 0.05. For *p* < 0.05, >1.3-fold for WT/KO, green; >1.3-fold for KO/WT; red, the others, black. *B*, Gene ontology analysis for the RNA-sequencing analysis. *p*-values obtained are shown as bar graphs. *C*, Mean difference (log2) in the expression levels in *Psg* (pregnancy-specific glycoprotein) and *Prl* (prolactin) gene families. *D*, RNA-sequence output with four replicate assays were averaged, compared between the genotypes, and plotted. Each plot indicates the respective genes in the family. Arrow indicates *Psg22*, which is mainly expressed in the early stage of pregnancy in normal mice.

### Aberrant endoreplication signatures in *Pparγ1*-KO placentas

Deletion of genes for canonical E2Fs (*E2f1-3*) promotes endoreplication (Chen et al., 2012). In contrast, when the genes for atypical E2Fs are deleted, endoreplication does not occur. We observed that sinusoidal TGCs had endoreplication signatures in the *Pparγ1*-KO labyrinth (Fig. 7A). In endoreplication, G_1_/S-phase is repeated with uncoupling of the replication–mitosis cell cycle. For these anomalous cell cycles, E2F transcription factors play a pivotal role (Li et al., 2008; Chen et al., 2012; Pandit et al., 2012); among these, canonical E2F1-3 mediates cell cycle exit, and atypical E2F7 and E2F8 drive endocycling. RT-qPCR analysis of labyrinth samples showed that the genes for the atypical type were increased *(E2f7* and *E2f8;* 3.5- and 18.8-fold, respectively) among *E2f* genes (Fig. 7A).

**Figure 7.**
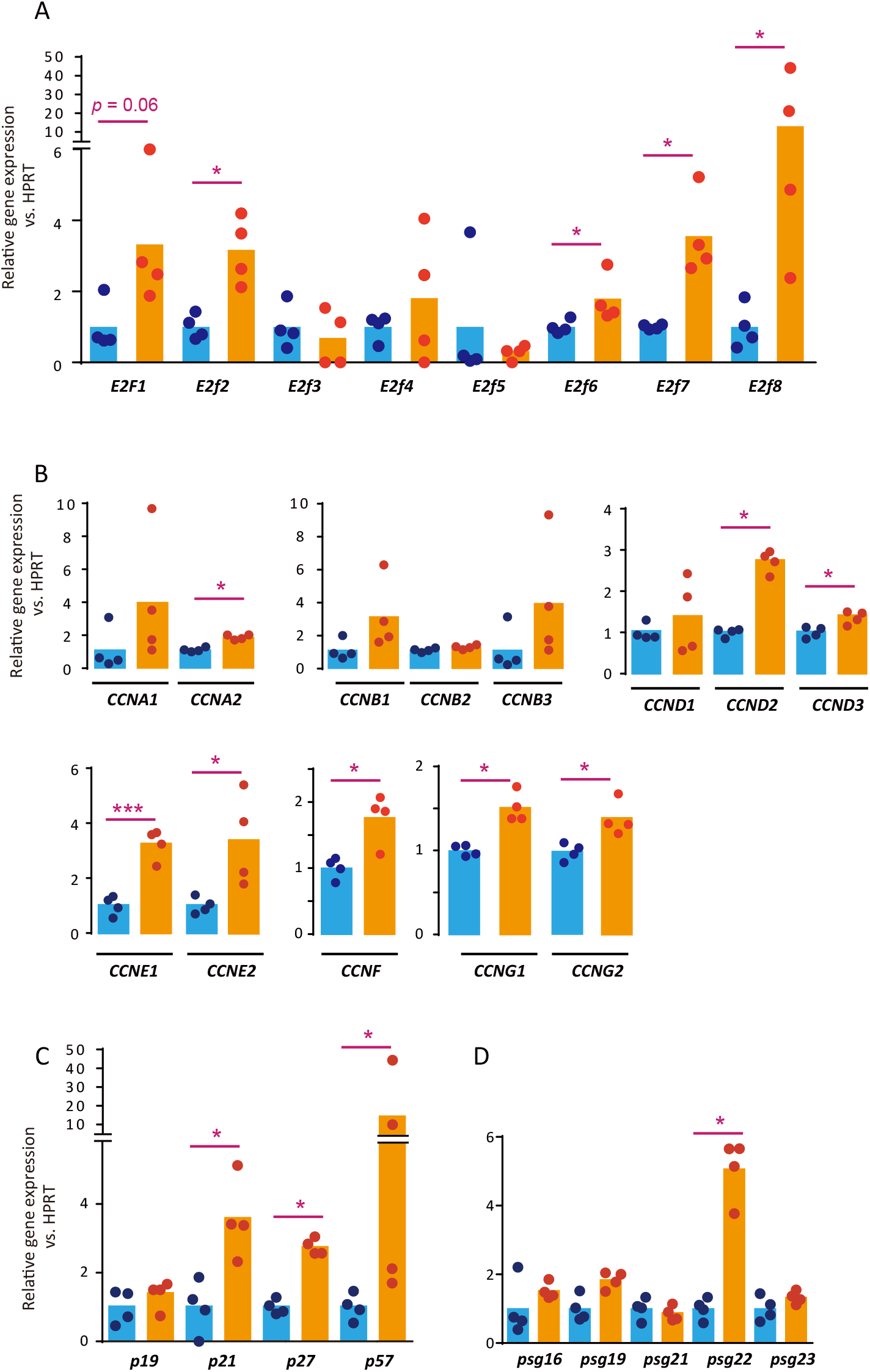
Quantitative RT-qPCR analysis of labyrinthine RNA for cell cycle-related gene expression levels (n = 4 per group). *p*-values were obtained using the Mann–Whitney U-test. *, *p* < 0.05; ***, *p* < 0.0001. *A*, genes encoding E2F family proteins; *B*, genes encoding cyclins; *C*, genes encoding cyclin-dependent kinase inhibitors; *D*, genes encoding pregnancy-specific glycoproteins.

Expression of cyclins is also a major determinant of cell cycle staging (Cuitiño et al., 2019). For S-phase, cyclin D and E types are increased. In endocycling, cyclins D (Soonpaa et al., 1997; Zimmet et al., 1997) and E (Geng et al., 2003; Parisi et al., 2003) play key roles. RT-qPCR analysis also confirmed that the gene expression levels for cyclin D and E types were greater in KO than the WT counterparts (Fig. 7B).

We also found that the other up-regulated genes (Fig. 7C), of which the products are involved in the endocycle: p57 inhibits cyclin-dependent kinase 1 that coordinates entry into mitosis; p21 inhibits the check-point protein kinase CHK1 (for endoreplication, see Ullah et al., 2008; Zielke et al., 2013); and p27 overlaps functionally with p21.

### Expression of gene families for *Psg* and *Prl* are dysregulated

In becoming polyploid, TGCs are metabolically strenuous, producing a variety of signaling molecules. The dysregulated cellular processes of sinusoidal TGCs may cause this production to deteriorate. With the RNA-sequencing conducted above, we found alterations in gene expression levels in *Pparγ1-KO* placentas. The expression levels of most genes related to pregnancy-specific glycoproteins (PSGs) were decreased (Figs. 6C-D). We quantified five PSG-coding genes in the labyrinth to find that *psg22* alone was increased approximately 5-fold in *Pparγ1-KO* placentas (Fig. 7D). *Psg22* was reported to be expressed predominantly in the early stage of TGCs during pregnancy (Williams et al., 2015), which might represent an immature state in sinusoidal TGCs. In addition, the expression of prolactin-related genes (*Prl*s) was dysregulated (Wynne et al., 2006) (Figs. 6C-D). *Muc-1* is a PPARγ-target gene for mucin, which is produced at the apical surface of the maternal blood sinuses in the labyrinth (Shalom-Barak et al., 2004), the expression of which was decreased by about a half in *Pparγ1-KO* placentas (Fig. 6D, bottom). These findings suggested that unbalanced trophoblast differentiation or cellular activity in *Pparγ1*-KO placentas underlies the eventual fetal growth restriction.

## Discussion

PPARγ is one of the key molecules in embryonic development, but its exact function is still elusive. The protein is abundant in the late stage of pregnancy in the conceptus. In the present study, by deleting *Pparγ1*, one of the PPARγ1-coding genes, we showed that the PPARγ1 protein is likely to be associated with sinusoidal TGC maturation, most likely in the endoreplication termination stage. The deletion resulted in enlargement of the nuclei of sinusoidal TGCs and disorganized placements, attenuating the development of feto–maternal interface architecture. *Pparγ1-KO* fetuses showed intrauterine growth restriction from 15.5 dpc, likely due to poor capillary formation in the labyrinth mainly without the appropriate functions of sinusoidal TGCs.

With the *Pparγ1*-specific deletion, the placentas developed normally in size and gross structure, which enabled us to examine the vasculature and cell morphology in detail. PPARγ has been implicated in TGC differentiation in mice and humans (Tarrade et al., 2001; Parast et al., 2009). In the present study, we observed sinusoidal TGC maturation impairment without *Pparγ1*, resulting in the deterioration of the feto–maternal boundaries, which was essentially consistent with the observations previously with *Pparγ*-full KO (Barak et al., 1999; Kubota et al., 1999). We also found similar and milder compromised labyrinthine structures in *Pparγ1sv-KO* placentas (Fig. S10). Thus, the degree of the deterioration presumably depends on the gene dosage of PPARγ1.

Polyploidization is suggested to increase cell size, genetic diversity, and protein production (Pandit et al., 2013). To initiate and drive the endocycle, coordinated interplays with various molecules are required (Sakaue-Sawano et al., 2013). A part of this process has been revealed, but the termination mechanisms have never been addressed. A number of studies showed that PPARγ agonists arrest the cell cycle, thereby inhibiting inhibit cell growth *in vitro* and in cancer cells inoculated into experimental animals (Law et al., 1996; Altiok et al., 1997; Motomura et al., 2000; Wakino et al., 2000; Itami et al., 2001; Koga et al., 2001; Bruemmer et al., 2003; Dios et al., 2003; Srivastava et al., 2014). Thus, PPARγ agonists are candidates for cancer therapy (Gou et al., 2017). With the placental transcript analyses, we showed that the deletion altered the expression patterns of E2Fs and cyclins, which are determinants for cell cycle staging (Cuitiño et al., 2019). The altered gene expression levels indicated promoted endoreplication in the KO, which is the first evidence for association of the cell cycle with PPARγ1 *in vivo*. Considering that PPARγ mediates terminal differentiation in a variety of cells, we speculated that without PPARγ1 protein sinusoidal TGCs are unable to regulate termination appropriately, preventing the cells from maturing. It is still to be determined what kinds of genes are targeted by PPARγ to coordinate or modulate the cell cycle.

The surface of the maternal circulation in the labyrinth provides sufficient nutrient exchange for the rapid growth of fetuses, especially in the late stage of pregnancy. The estimated surface of the maternal blood sinuses increases more than 3-fold from E14.5 to E16.5 in mice (Coan et al., 2004), and the fetuses grow more than 3-fold (approximately 0.4 g to 1.4 g) from 15.5 dpc to term (Fig. 2H) (Spangenberg et al., 2014). PPARγ1 is thought to mediate the placental adaptations that occur from E14.5 to E16.5 (Coan et al., 2004). This period is also when *Pparγ1* and *Pparγ1sv* expression levels peaked (Fig. 1) and the growth restriction of *Pparγ1*-KO fetuses started (Figs. 1G-H). Thus, abundant PPARγ1 around 15.5 dpc plays a crucial role in the functional adaptation of the placenta. This possibility is still a speculation because of the limited molecular evidence in the present study. In addition, it remains to be examined what are the differences between early death with *Pparγ1-full* KO and the growth restriction in this study.

The results of the present study suggested that PPARγ mediates endocycle exit, and its absence allows sinusoidal TGCs to maintain the endocycle but prevents them from maturing, thereby leading to fetal death in the end. The findings and materials from this study provide opportunities to reveal how PPARγ is involved in development via cell cycle regulatory functions.

## Materials and Methods

Table S3 provides detailed information on the materials, instruments, animals, and software used in this study.

### Targeting vectors

We constructed a *Pparγ* targeting vector consisting of a Neo encoding cassette, FRT-PGK-gb2-neo-FRTloxP, for exon A1 and exon C deletions. A positive bacterial artificial chromosome clone was used as a template to amplify genomic DNA fragments of *Pparγ*. An approximately 13.9-kb genomic fragment containing exons A1 and C was inserted into the targeting vector. The resulting targeting vector (Fig. S3) was linearized before electroporation.

### Generation of mouse strains with *Pparγ1*^+/-^ and *Pparγ1sv*^+/-^

Mouse strains with deletions of either exon A1 or exon C were generated using an embryonic stem cell method. First, we electroporated murine ES cells generated from C57BL (Kitayama et al., 2001) with the targeting vectors (Fig. S3A) and cultured homologous recombinant clones at TransGenic Inc. (Fukuoka, Japan). The resulting exon A1-floxed mouse line was intercrossed with B6;D2-Tg(CAG-Flp)18Imeg to generate the *Pparγ1*-KO allele (Figs. S1, S3A). Similarly, an exon C-floxed mouse line was intercrossed with B6;D2-Tg(CAG-Flp)18Imeg to delete the Neo cassette and successively with B6;CBA-Tg(CAG-Cre)47Imeg to generate the *Pparγ1sv*-KO allele (Figs. S1, S3A). *Pparγ1*^+/-^ and *Pparγ1sv*^+/-^ mice were backcrossed with male C57BL/6J mice 8 and 5 times, respectively.

### Mice

Adult male C57BL/6J mice were purchased from a domestic provider. All the mice were housed in a pathogen-free facility with food and water ad libitum in accordance with guidelines set by the Animal Care and Use Committee, Saitama Medical University.

### Timed mating and embryo processing

Mouse estrous cycles were monitored by examining vaginal smears with May-Grüwald Giemsa staining (Cora et al., 2015). Heterozygous female mice (5 weeks to 6 months old) were weighed, crossed with heterozygous males (8 weeks to 8 months old) at 3 PM. The next day, we separated female mice with a copulatory plug at 9 AM. We set the next morning as 0.5 dpc. The conceptuses were excised at the indicated timepoints under deep anesthesia with isoflurane, and the dams were sacrificed after the removal. Gross morphology of the conceptuses was examined if necessary with the use of a stereomicroscope.

### RT-qPCR

Total RNA was isolated using SV Total RNA Isolation System (Promega) from the placentas and reverse-transcribed (SuperScript IV VILO Master Mix, ThermoFisher). Quantitative expression analysis was performed with SYBR^®^ green technology. The expression levels of *Pparγ1*, which includes exon A1, *Pparγ1sv*, which includes exon C, *Pparγ2*, and *18S* ribosomal RNA were quantified using linearized full length cDNAs as standards (Virtue et al., 2010; Takenaka et al., 2013). For comparison using the ΔCT method with calibrated PCR efficiency, the levels were normalized to that of *HPRT* to minimize sample variance. The levels were further normalized to those of the control groups. The variables were compared using the Mann–Whitney *U*-test. The primers used are listed in Table S4.

### Western blotting

Proteins were separated on 12% polyacrylamide gels and transferred to polyvinylidene difluoride membranes. An anti-PPARγ polyclonal antibody and horseradish peroxidase-conjugated anti-rabbit IgG antibody were used as the primary and secondary antibodies, respectively. Bands were visualized using a chemiluminescence kit with ChemiDoc^™^ MP system. MagicMark^™^, a chemiluminescent protein size marker, was used. As a PPARγ positive control, a nuclear fraction of adipocyte-like differentiated 3T3-L1 cells (Takenaka et al., 2013) was loaded together with the samples. We adjusted the signal levels when necessary using Photoshop CS5 software to enhance the visibility of bands. The signal intensity of each band was obtained using ImageJ. We assumed that the protein abundance loaded onto the gel was consistent, as shown in Fig. S13B.

### RNA-sequence

Procedures for RNA-sequencing and data analysis are shown in the Supplementary Materials and Methods in detail. The sequences read can be found in BioProject with accession number PRJNA626683.

### Histology

Histological analysis was performed on three different placentas at 15.5 dpc obtained from 2–3 different mice. Reagents for histological staining were obtained from Muto Pure Chemicals (Tokyo, Japan). Uteruses were excised from anesthetized pregnant mice, submerged in 4% paraformaldehyde, and cut between implantation sites. The conceptuses were dissected as described in (David et al., 2006) and immersed in 4% paraformaldehyde overnight, dehydrated in a gradient alcohol series, and embedded in paraffin. Thin sections (4 μm) were cut and stained the H&E for general morphology, Masson trichrome for blood vessel visibility, PAS to identify the junctional zones, and Feulgen to estimate nuclear abundance. For the nuclear abundance estimation, the green channel of the images was extracted, the pixel intensity was inverted, and the density of chromosomes was quantified. Background pixel density (a tissue negative area) was subtracted, and divided by the area to normalize nuclear density.

TGC nuclear volume was calculated using the estimated radius [(long radius + short radius)/2]. Images were obtained with the use of a microscope and analyzed with ImageJ in accordance with the method of Schindelin et al. (Schindelin et al., 2012).

### Immunohistochemistry

Deparaffinized and hydrated thin sections were antigen-retrieved by heating at 60°C in 1 mmol/l EDTA (pH 8.0) overnight. The sections were blocked by 3% H_2_O_2_ for endogenous peroxidase activity. We used a Vector^®^ M.O.M.^™^ Immunodetection Kit for the anti-PPARγ mouse monoclonal antibody. We used 0.5 mg/ml 3,3-diaminobenzidine to detect peroxidase activity. The sections were counterstained with methyl green (Muto Pure Chemicals) where appropriate. Area for each puncta was quantified using ImageJ.

We also stained placental sections for MCT-1 with anti-MCT-1 IgY and followed by Alexa Fluor 488-conjugated goat anti-chicken IgY. DAPI to stain the nuclei.

### Transmission electron microscopy

After heterozygous crossing, female mice were sacrificed at 15.5 dpc, and the labyrinthine zones were dissected and fixed in 2.5% glutaraldehyde for 1 day and postfixed in 1% OsO4 solution buffered with Sorenson’s phosphate at pH 7.4. Then, the samples were rapidly dehydrated in alcohol and embedded in Epon 812. Thin sections of representative areas were examined using an electron microscope (JEM-1400 Transmission Electron Microscope, JEOL Ltd., Tokyo, Japan).

### Statistical analysis

The data are shown as means with individual plots unless otherwise indicated. Nonparametric data are shown as box plots with a median, box (25–75th percentile), and bar (5–95th percentile) with outliers. The data mainly derive from three genotypes (WT, Het, and KO). They were compared using one-way ANOVA when the data were considered to be parametric. Individual differences were evaluated using Student’s *t* test or the Dunnett test if the ANOVA was significant and the test was necessary. If the data showed nonparametric distribution, the data were compared using the Steel–Dwass test or Mann–Whitney U test. Distribution of genotypes yielded from heterozygous crosses were examined using Hardy–Weinberg’s law test.

## Acknowledgements

The authors greatly thank Ms Sato Sawako, Ms Yuka Nakano, and Ms Reiko Inomata for their technical assistance, and Dr Norihiro Kotani for providing reagents used in the fluorescence immunohistochemistry. We are also grateful to members of the Biomedical Research Center, Saitama Medical University, for their help with the present work, especially Ms Kumiko Komatsu for help in transmission electron microscopy experiments, Ms Noriko Murai for technical assistance in scanning electron microscopy experiments, Dr Susumu Ohshima for technical assistance with microscopy, and Ms Motoko Inomata for assistance with the histological analyses. This study was supported by JSPS KAKENHI grants to I.I. (Grant numbers 26461367, 25460299, and 17K09866).

## Author contributions

**T.N.**: Data curation, investigation, writing – original draft preparation, writing – review and editing, visualization, project administration **H.A**.: Investigation, resources **M.H.**: Software, formal analysis, investigation **Y.T.**: Methodology, writing – review and editing, visualization **K.F.**: Investigation, validation **H.S.**: Validation **H.K.**: Investigation, validation **T.K.**: Investigation **A.O.**: Software, supervision **T.M.**: Supervision **A.K.**: Conceptualization, supervision **I.I.**: Conceptualization, resources, supervision, funding acquisition.

