## Supplementary materials for "Targeted disruption of *Pparγ1* promotes trophoblast endoreplication in the murine placenta"

#### Supplementary Methods

*Mice*

Pregnant C57BL/6J mice were purchased from Clea (Tokyo, Japan) at 7 or 10 days post-coitum (dpc). The mice were sacrificed at 10, 12, 14, 16, and 18 dpc under deep anesthesia with isoflurane to obtain the conceptuses. The placentas and fetuses were separated, except for 10 dpc, and they were frozen immediately. The tissues were used to estimate gene expression of *Pparγ1*, *Pparγ1sv*, and *Pparγ2* with reverse transcript quantitative PCR (RT-qPCR) analysis.

*Genotyping*

For PCR genotyping, a KOD FX Neo kit (TOYOBO life Science, Tokyo, Japan) was used. DNA was obtained from proteinase K-digested tail and yolk sac biopsies from adult mice and fetuses, respectively. Primers used were as follows: F1: 5'-ATT CGC CTT CAT AAC ATT CT-3'; F2: 5'-TGG TCT GGC TGT GTT CTT GTA CTG-3'; F3: 5'-GTA ACT GAC AGC CTA ACC CT-3'; F4: 5'-TGT GCT CGA CGT TGT CAC TGA A-3'; R1: 5'-TGC TGC TCC AAA TGC TCG TAG TAT C-3'; R2: 5'-CCT CAG ACC GAT GTC CAT G-3'; R3: 5'-CGA GCC CCT CTC TAA ATC TGT-3'.

*Scanning electron microscopy*

Deparaffinized and hydrated thin placental sections were treated with TI blue (Nisshin EM Co., Ltd., Tokyo, Japan) and scanned using a Miniscope® TM3000.

*RNA-sequence*

The RNA library was prepared from high-quality RNA depleted of rRNA. Following adaptor ligation, the resulting DNA was amplified by PCR for 12 cycles and purified. Libraries were quantified and then used for cluster generation and sequencing. For sequencing details, see below.,

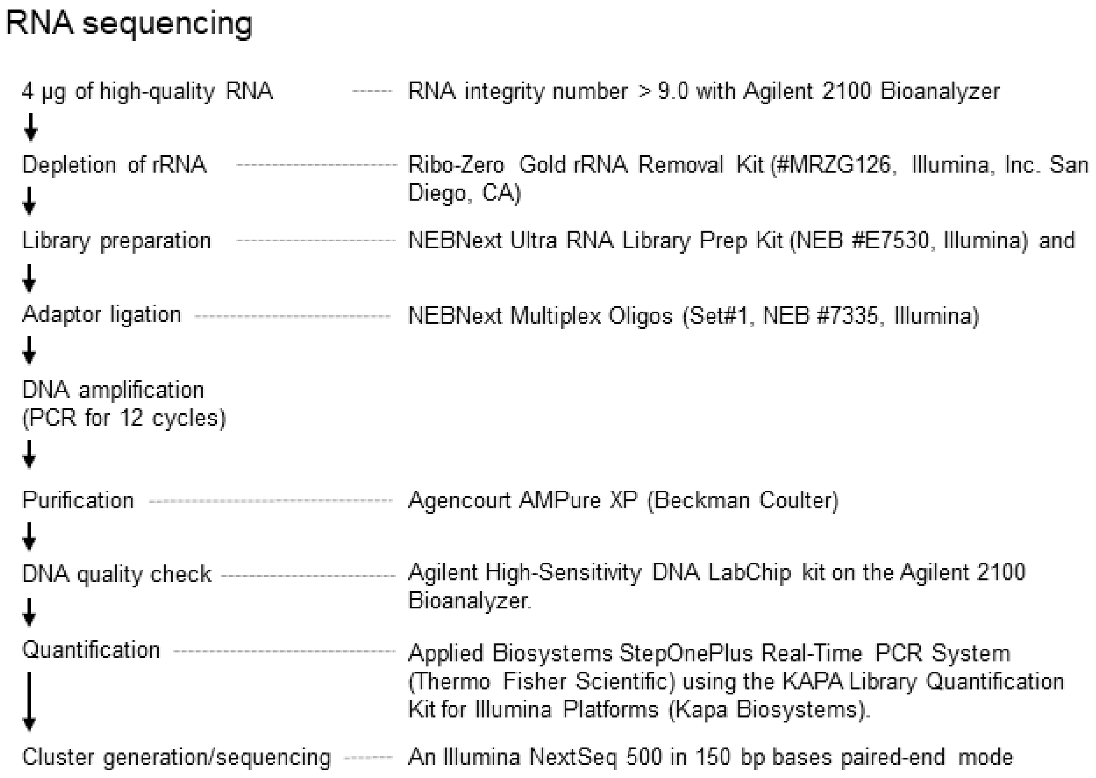


*Analysis of RNA-sequence data*

Low-quality bases were removed from the reads above, and the resulting trimmed sequencing reads were aggregated into a rRNA reference to remove rRNA reads. Then, the clean reads were mapped to the grcm38_snp_tran reference genome and sorted. Gene expression levels were measured with FPKM (fragments per kilobase of exon per million reads mapped) calculated using R package Ballgown. The readcounts were calculated by using HTseq. *P* values for the difference among genotypes were obtained using the edgeR package (https://bioconductor.org/packages/release/-bioc/html/edgeR.html). DAVID (https://david.ncifcrf.gov/) was used for GO analysis. $p < 0.05$ was considered to be significant in the analyses. For the analysis procedure details, see the flow chart on the next page.

*Analysis of RNA-sequence data* (*Continued*)

#### Data analysis

RNA-sequencing data

↓

Removal for low-quality bases ------- Phred quality score <20, FASTQ Quality Trimmer (FASTX Toolkit, http://hannonlab.cshl.edu/fastx_toolkit/)

↓

Removal for rRNA reads ------------ Aggregated into rRNA reference (Mouse rRNA reference BW000964) using Bowtie (http://bowtie-bio.sourceforge.net/index.shtml)

↓

Mapping ------------ The grcm38_snp_tran reference genome using HISAT2 (http://daehwankimlab.github.io/hisat2/) with default parameters

↓

Sorting ------------ SAMtools (http://samtools.sourceforge.net/)

↓

Assembling ------------ StringTie; the "- G" option for GRCm38_genes.gff and the 8 assembled transcript files (GTF format) were merged.

Readcount ------------ HTseq-count (https://htseq.readthedocs.io/en/release_0.11.1/ )

False discovery rate (FDR) ---- The edgeR package, an R-based tool within the Bioconductor project based on the featurecounts

↓

FPKM* reads mapped ------------ Ballgown bundled in R package (http://bioconductor.org/packages/release/bioc/html/ballgown.html)

*FPKM, Fragments per kilobase of exon per million

#### Supplementary figures

*Supplementary Figure 1. Exon structure for Pparγ-related genes*

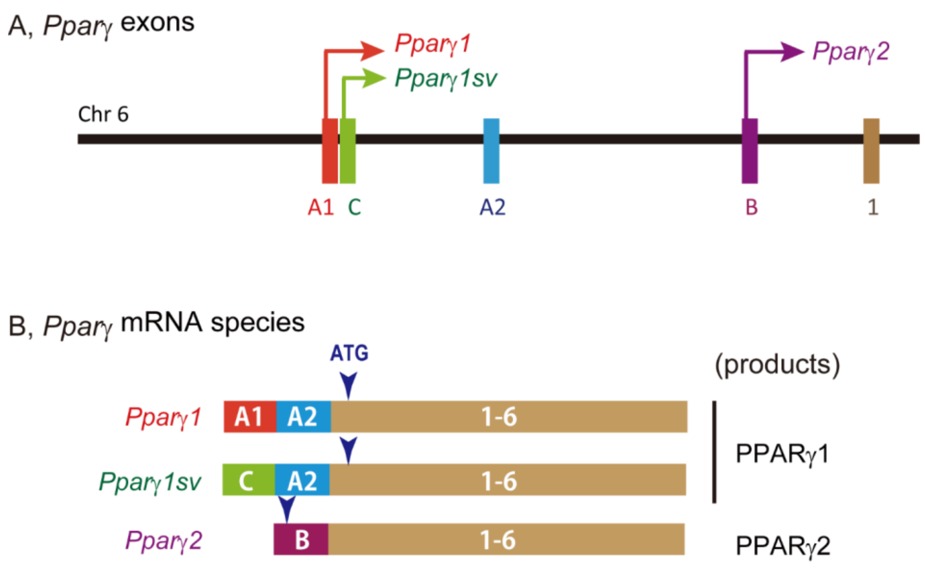


*A*, Structure of *Pparγ* exons on chromosome (chr) 6. *Pparγ1* and *Pparγ1sv* start at exon A1 and exon C, respectively. *B*, Resulting mRNA structures for *Pparγ* species. Exon A1 and exon C are specific transcripts for *Pparγ1* and *Pparγ1sv*, respectively. Modified from Takenaka Y, Inoue I, Nakano T, Shinoda Y, Ikeda M, Awata T, Katayama S (2013), A Novel Splicing Variant of Peroxisome Proliferator-Activated Receptor-γ (*Pparγ1sv)* Cooperatively Regulates Adipocyte Differentiation with *Pparγ2*. PLoS ONE 8:e65583 [This paper was published under a CC BY license].

*Supplementary Figure 2. CAGE data for Pparγ1 and Pparγ1sv in the placenta*

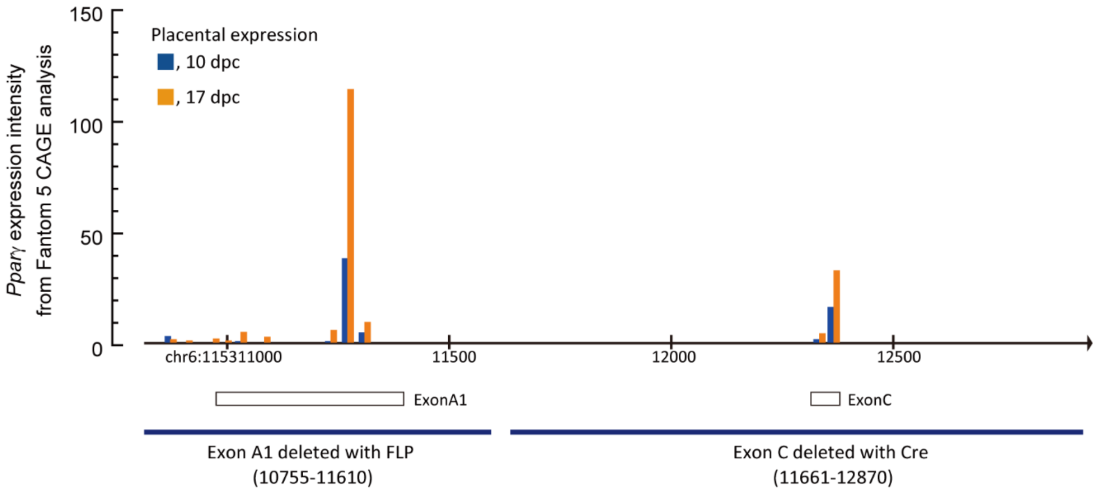


The X-axis shows the genome region that includes exon A1 and exon C on chromosome (chr) 6 refers the mm9 database. The Y-axis shows the transcription start sites and their density for *Ppar*γ expression at 10 dpc (*yellow*) and 17 dpc (*blue*) in mice obtained at the FANTOM5 mouse promoterome view (http://fantom.gsc.riken.jp/zenbu/ accessed in July, 2019). The boxes under the X-axis indicate the sites of the exons. The blue bars at the bottom indicate sites that were genetically deleted for the development of transgenic mouse lines in the present study.

*Supplementary Figure 3. Exon A1- or exon C-specific deletion in the mouse genome*

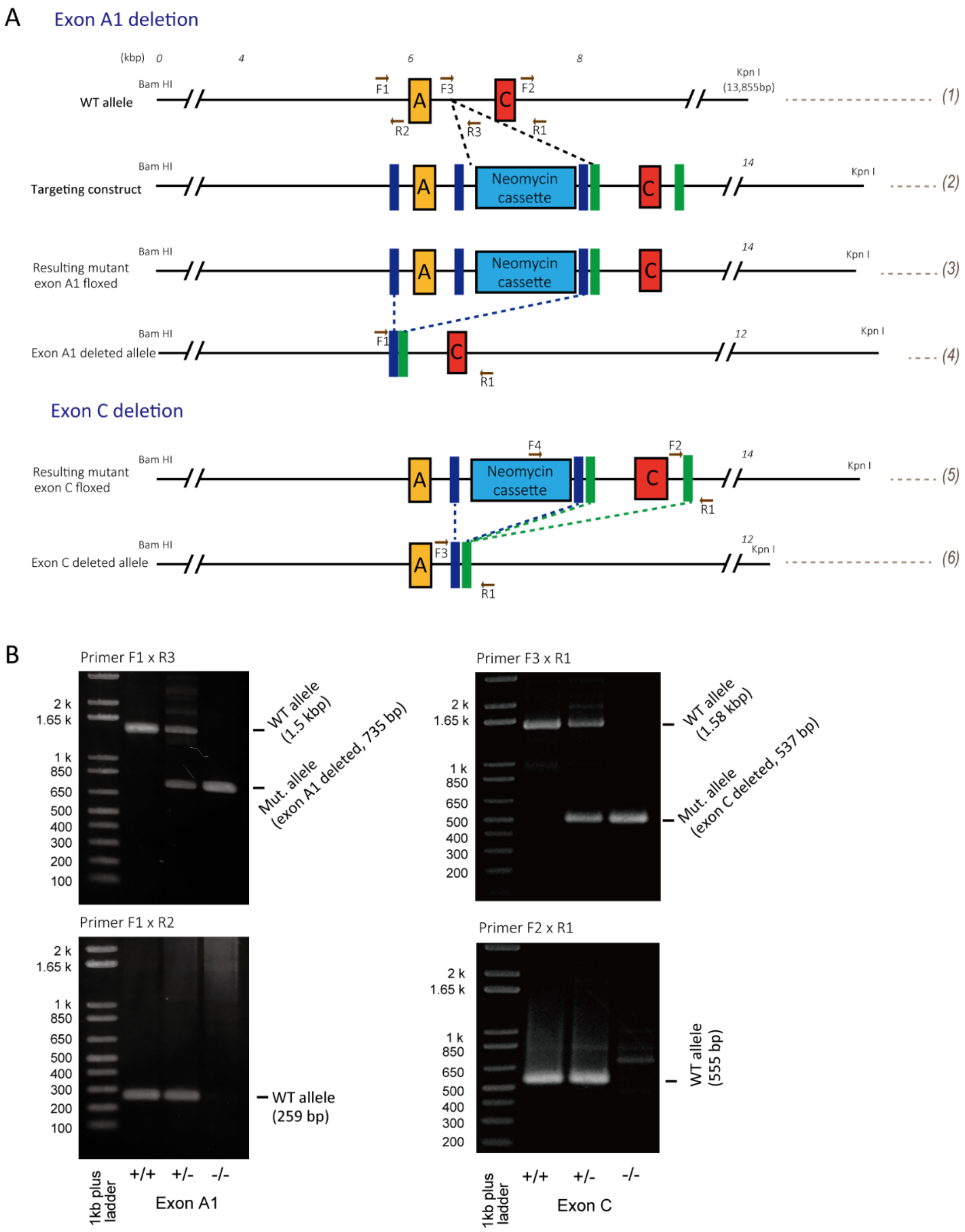


*A,* Exon A1 and exon C deletion procedures using gene targeting. *(1) WT* allele represents the genomic region containing exon A1 and exon C. *Arrows*, the sites used for priming in PCR. For primers, see Table S2. *(2) Targeting construct* used for homologous recombination with the WT allele. Blue bars, FRT sites; green bars, LoxP sites. *(3)* The resulting mutant allele was floxed for exon A1. Mice with the allele *(3)* were crossed to those with FLP to produce an allele lacking exon A1 *(4)*. *(5,6)* Exon C deletion procedure. *(5)* The resulting mutant allele was floxed for exon C. Mice with the allele *(5)* were crossed to those with Cre to produce allele *(6)* lacking exon C. *B*, Genotyping of resultant mutants with PCR. F1 × R3 and F1 × R2 primer pairs were used for exon A1 deletion allele genotyping. F3 × R1 and F2 × R1 were used for exon C deletion.

*Supplementary Figure 4. Pparγ1sv-KO male mice had lower body weight.*

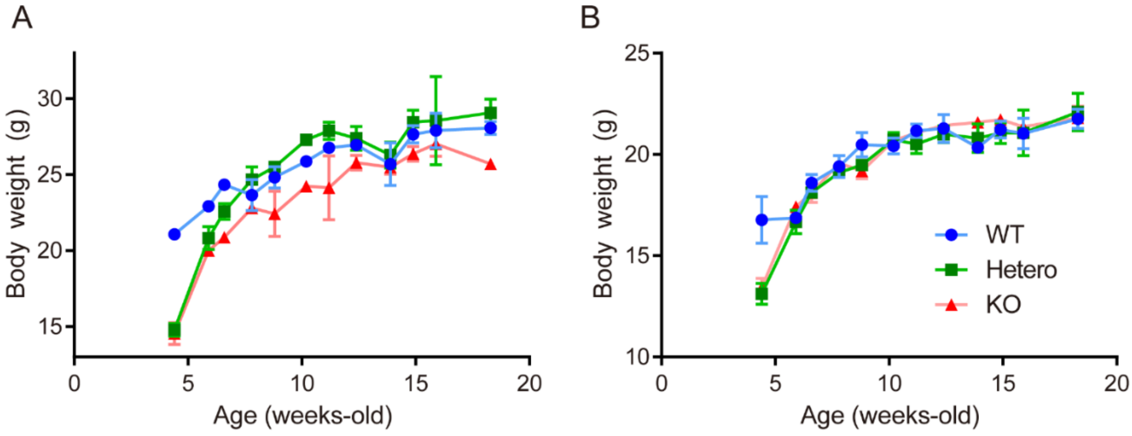


***Two-way ANOVA analysis for A and B***

| Male | | | Female | | |
|---|---|---|---|---|---|
| Factors | *F* (DFn, DFd) | *p*-value | Factors | *F* (DFn, DFd) | *p*-value |
| Interaction | *F* (22, 142) = 2.120 | 0.005 | Interaction | *F* (22, 158) = 1.721 | 0.03 |
| Age | *F* (11, 142) = 47.83 | < 0.0001 | Age | *F* (11, 158) = 60.14 | < 0.0001 |
| Genotype | *F* (2, 142) = 24.82 | < 0.0001 | Genotype | *F* (2, 158) = 2.709 | 0.07 |

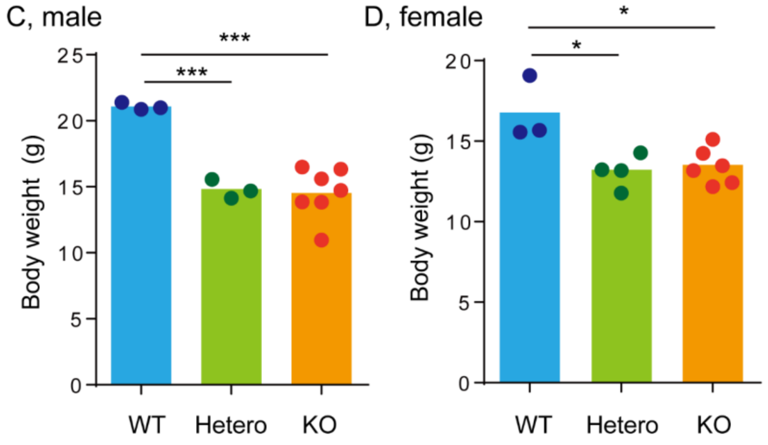


Body weights of *Pparγ1sv*-KO and -Het mice were smaller than the WT counterparts at post-weaning. *A*, In male mice, body weight of KO tended to be smaller than the other counterparts over the time points measured (for genotype, $p < 0.0001$ by two-way ANOVA, see inset). *B*, In female mice, such significant differences were not observed ($p = 0.07$). Female mice showed similar body weight between the genotypes from 5 to 18 weeks old. Body weight was measured occasionally for 13 WT, 16 Het, and 12 KO male mice; for 14 WT, 17 Het, 15 KO female mice. The number of data points ranges from 1 to 13 for each plot. Data are shown as mean and SEM. The data used were not sequential; thus, we did not statistically analyze the differences between the groups. *C* and *D*, Comparison of body weights from 4 to 5 weeks old showed significantly lower body weights in mice with exon $C^{+/-}$ and exon $C^{-/-}$. Statistical significance was obtained using Students-t test after one-way ANOVA analyses (for male, $F$ (2, 10) = 20.89, $p = 0.0003$; for female, $F$ (2, 10) = 8.070, $p = 0.008$). *, $p < 0.05$; ****, $p < 0.0001$.

*Supplementary Figure 5. Preliminary examination to detect PPARγ protein using western blotting*

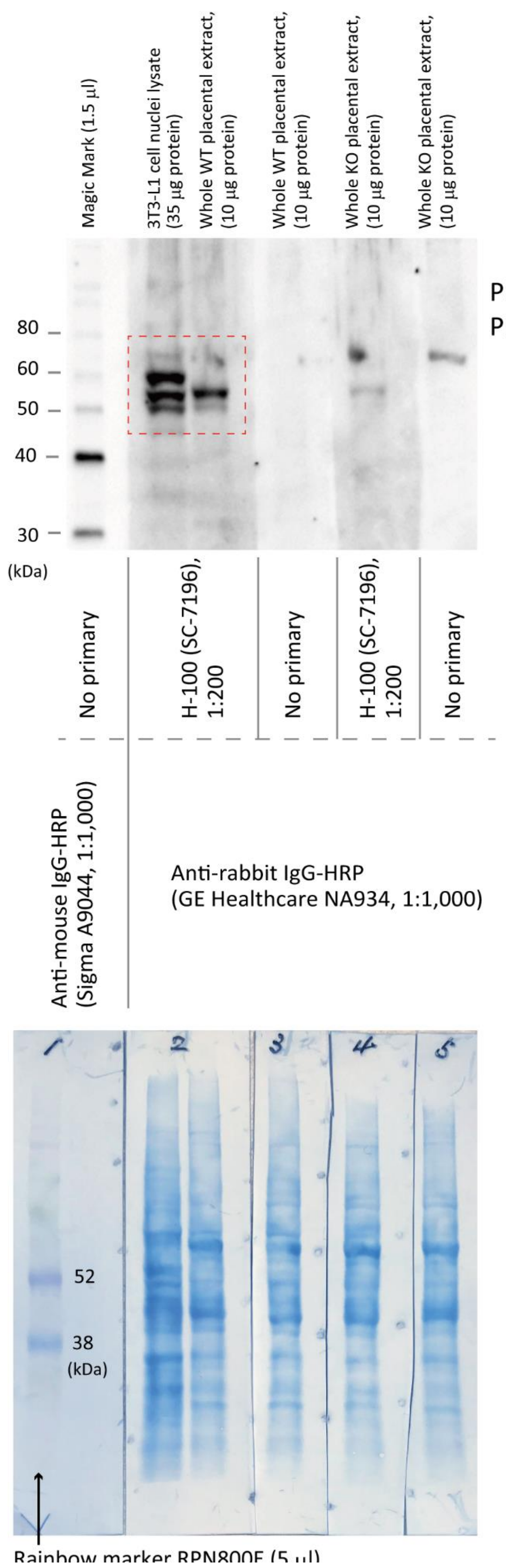


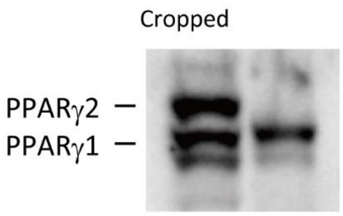


*Upper*, Adipocyte differentiated 3T3-L1 cells were used as a positive control for PPARγ. Whole protein extracts from a WT placenta was loaded in the neighboring lane, showing that PPARγ1, but not PPARγ2, protein was present (also see the cropped picture in the *right*). The lane with no primary antibody had no bands. The extract from a KO-placenta had a weak PPARγ1 protein band.

*Bottom*, Proteins transferred to a membrane and stained with CBB.

*Supplementary Figure 6. Mice with Pparγ1$^{+/-}$ develop and grow normally.*

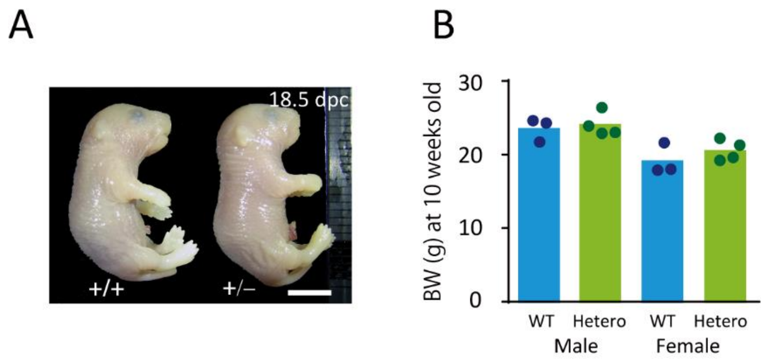


*A*, No developmental retardation was observed in *Pparγ1*$^{+/-}$ mice.

*B*, No difference between WT and Het in body weight at 10 weeks old.

*Supplementary Figure 7. PPARγ expression in the labyrinth at the 15.5 dpc*

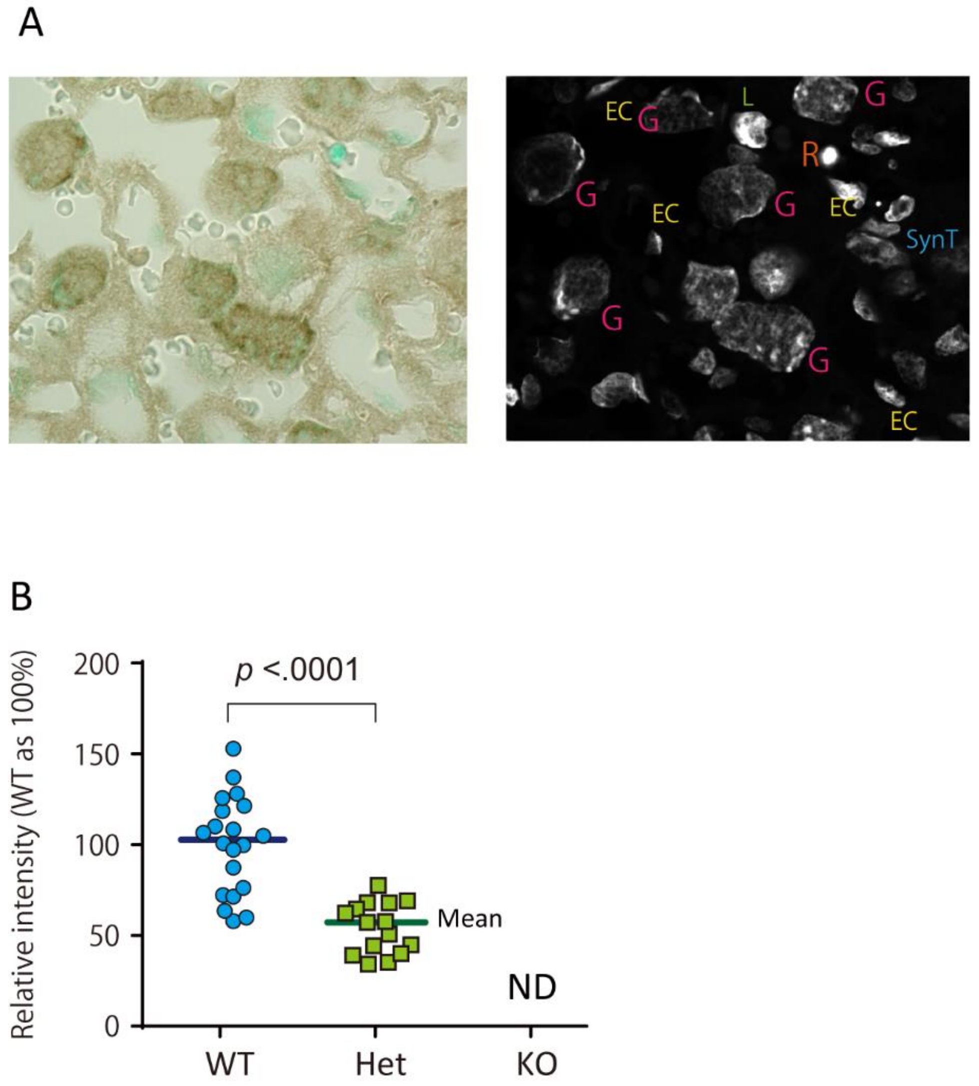


*A*, *Pparγ1*-WT staining with PPARγ (DAB) and nuclei (methylene blue). *Left*, bright field microscopy; *Right*, methylene blue fluorescence image. G, trophoblast giant cells; EC, endothelial cells; R, red blood cell; SynT, syncytiotrophoblast. *B*, Quantification of DAB intensity. *Pparγ1*-KO placentas were not examined because apparent staining was not observed. WT and heterozygotes were compared using Student's *t*-test. ND, not determined. Bars indicate means.

*Supplementary Figure 8. PPARγ expression in other trophoblast giant cells.*

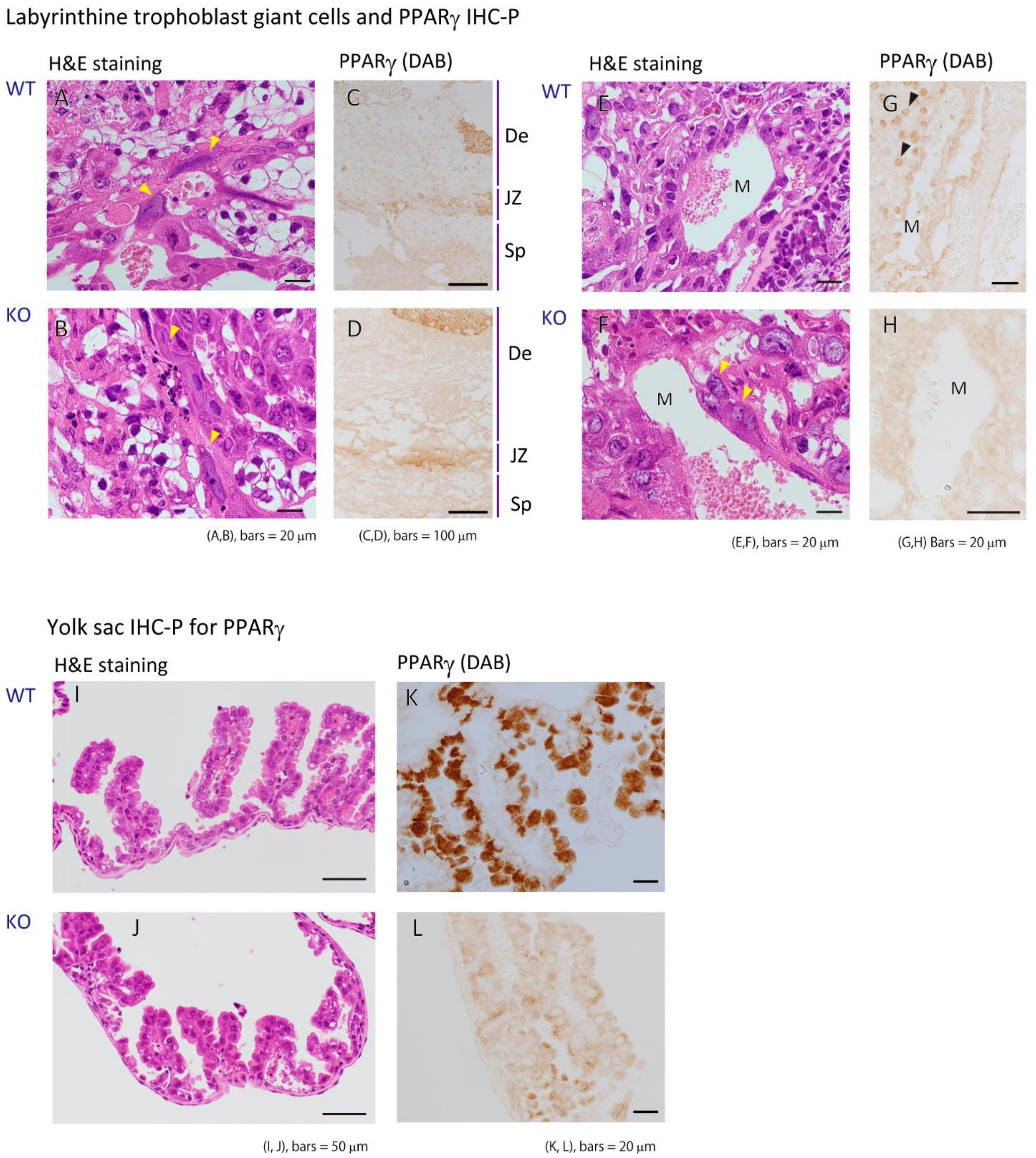


*A-H*, No apparent PPARγ-immunostaining in parietal (*A-D*) and maternal blood canal-associated (*E-H*) TGCs. Yellow arrow heads in panels *A*, *B*, and *F* indicate TGCs. Black arrow heads in panel G indicate the presence of sinusoidal TGCs in the labyrinth. De, decidua; JZ, junctional zone; Sp, spongiotrophoblast. *I-L*, Histological analyses of yolk sac at 15.5 dpc. *I* and *J*, H&E staining for the yolk sacs. *K* and *L*, PPARγ immunostaining for cuboidal epithelial cells in the yolk sac (*K*). Intensive staining can be seen only in WT, but not KO (*L*). For the discrimination of TGC species, see “Hu D, Cross JC (2010), Development and function of trophoblast giant cells in the rodent placenta. *Int J Dev Biol* 54:341-354.”

*Supplementary Figure 9. Scanning electron microscopy of the labyrinth zone*

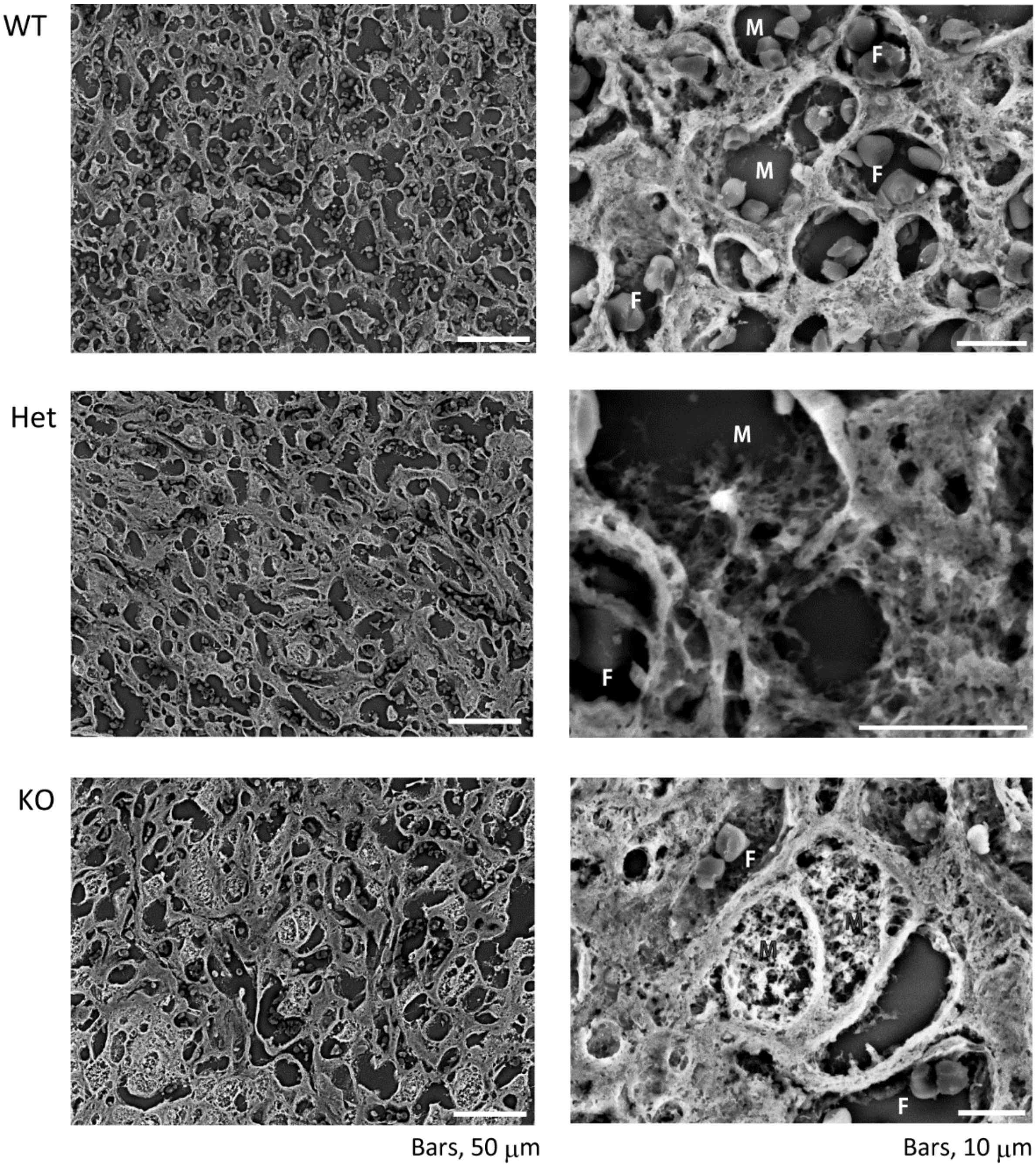


SEM shows that maternal blood sinuses (M) are round and have a smooth surface. Deletion of the *Pparγ1* gene made them squashed and enlarged. The surfaces were coarse, especially in the KO. F, fetal blood capillary.

*Supplementary Figure 10.*

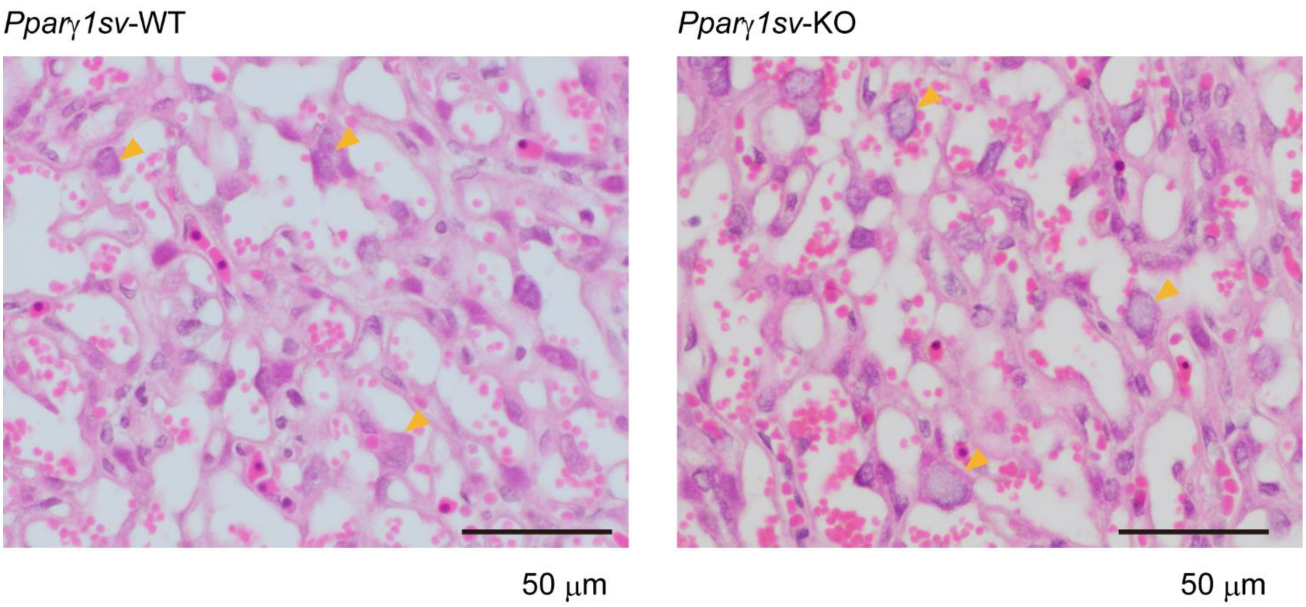


H&E staining of labyrinth in placentas carrying male embryos obtained from pregnant mice at 15.5 dpc. Right and left panels show *Pparγ1sv*-WT, and -KO labyrinths of the placentas, respectively. Arrowheads indicate the nuclei of sinusoidal trophoblast giant cells.

*Supplementary Figure 11. Transmission electron microscopy analysis of the labyrinth*

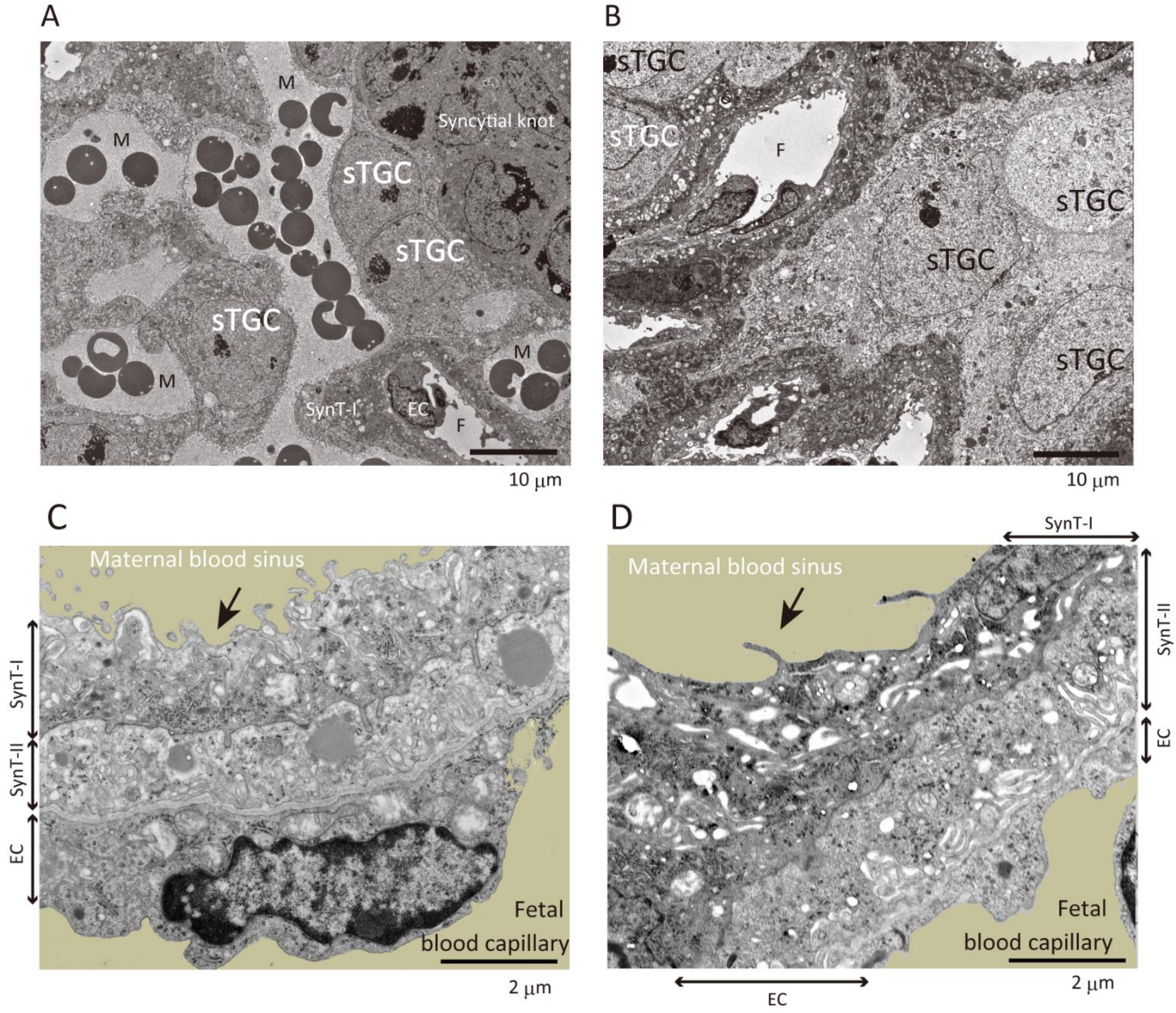


WT on the *left* (*A, C*), KO on the *right* (*B, D*). *A* and *B* show the spatial localization of sinusoidal TGCs (sTGC) in the section. Sinusoidal TGCs were not facing the maternal blood sinus in the KO (*B*). *C* and *D* show feto–maternal interfaces. Poor microvillus development to the KO maternal blood sinus was apparent, as indicated by arrows (*D*). TGC, trophoblast giant cells; M, maternal blood sinus; SynT, syncytiotrophoblast; EC, endothelial cell; F, fetal blood capillary. Bars in *A* and *B*, 10 μm; 2 μm in *C* and *D*.

*Supplementary Figure 12. Effect of Pparγ1-deletion on the placental gene expression*

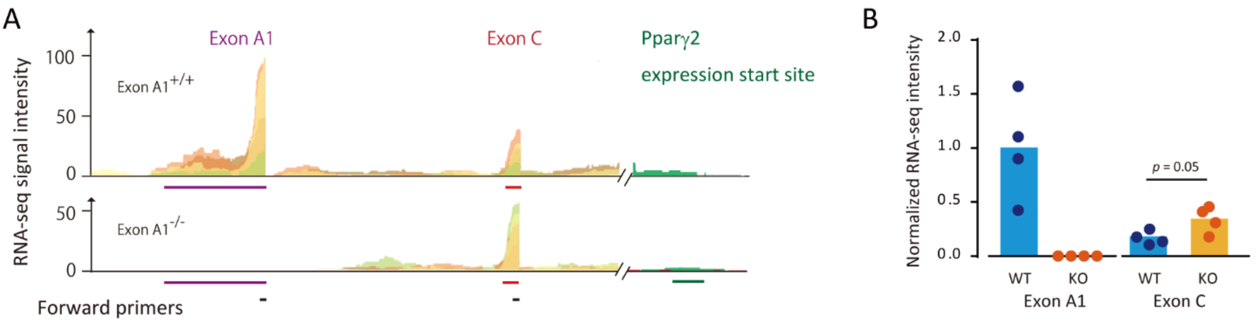


C

| GO analysis for placental RNA-sequencing output | |
|---|---|
| GO Term | P-Value |
| Up-regulated | |
| Nucleosome assembly | 8.0E-16 |
| DNA methylation on cytosine | 4.0E-14 |
| Negative regulation of megakaryocyte differentiation | 4.2E-06 |
| DNA-templated transcription, initiation | 6.1E-05 |
| Cell-cell adhesion | 2.6E-04 |
| Negative regulation of transcription from RNA polymerase II promoter | 6.4E-04 |
| Membrane raft assembly | 6.8E-04 |
| Phosphorylation | 9.9E-04 |
| Down-regulated | |
| Female pregnancy | 3.2E-10 |
| Inflammatory response | 1.4E-06 |
| Response to hypoxia | 2.8E-05 |
| Positive regulation of inflammatory response | 1.3E-04 |
| Positive regulation of angiogenesis | 1.3E-04 |
| Positive regulation of gene expression | 1.4E-04 |
| Positive regulation of cell migration | 2.3E-04 |
| Aging | 3.1E-04 |
| Axon guidance | 5.5E-04 |
| Proteolysis involved in cellular protein catabolic process | 6.9E-04 |
| Retrograde protein transport, ER to cytosol | 9.3E-04 |

*Supplementary Figure 12 (continued)*

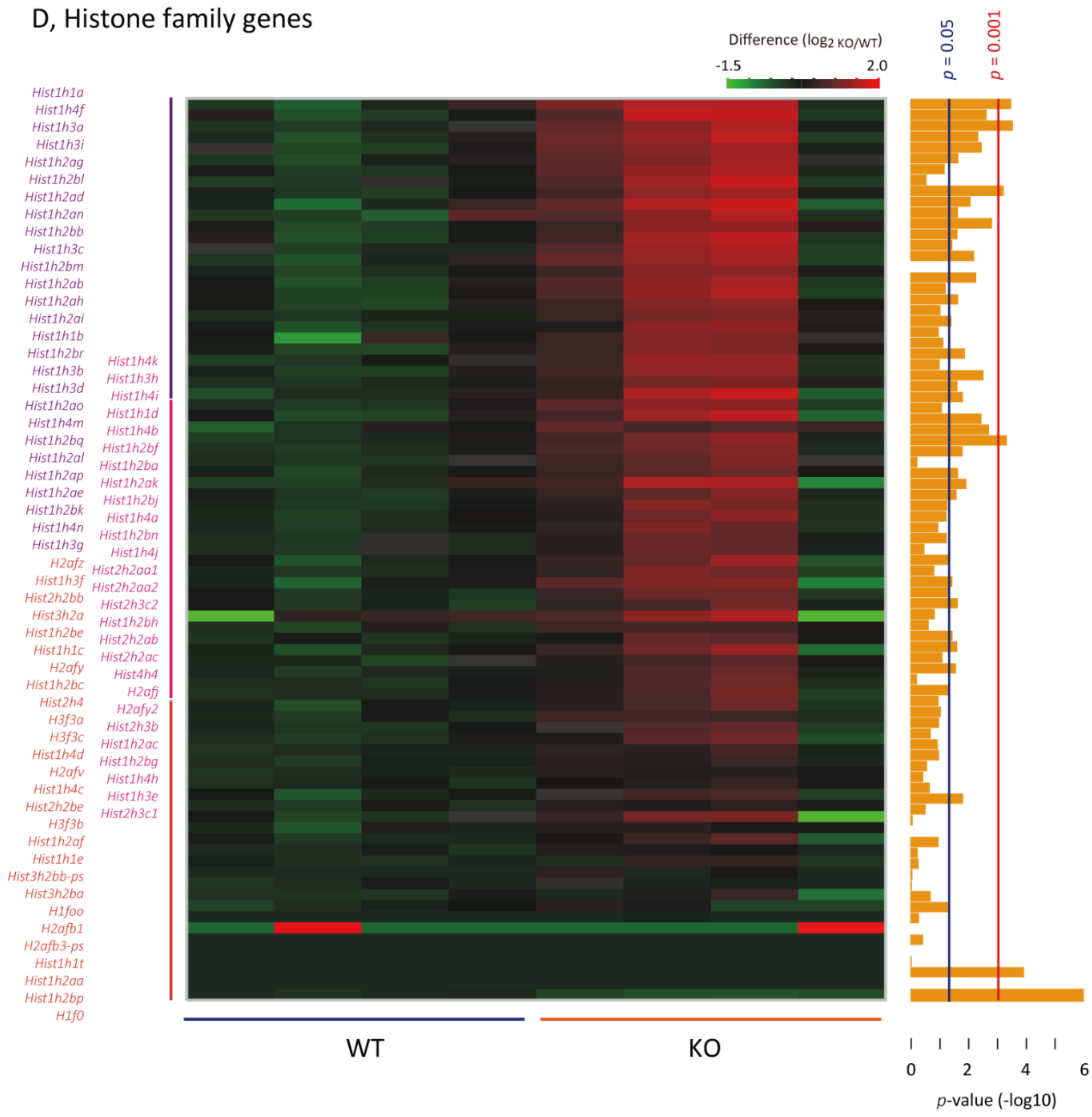


RNA-sequencing analysis reveals endoreduplication signature and pregnancy-related gene expression dysregulation

*A*, Gene expression of *Pparγ* species starting sites (X-axis) and the frequency (Y-axis) are visualized for four replications. Each first exon is indicated with the respective bars. *B*, Comparison of signal intensity using RNA-sequencing between WT and KO. *P* value was obtained using Student's *t*-test. *C*, Tabulated results of gene ontology analysis. *D*, Heatmap presentation for histone-related gene expressions by RNA-sequencing analysis of placentas at 15.5 dpc (n = 4 per genotypes). $Log_2$ fold changes are pseudocolored with indicated ranges. *p*-values are converted to common logarithm and shown as bar length on the *left* as $log_{10}$ *p*-values.

*Supplementary Figure 13*

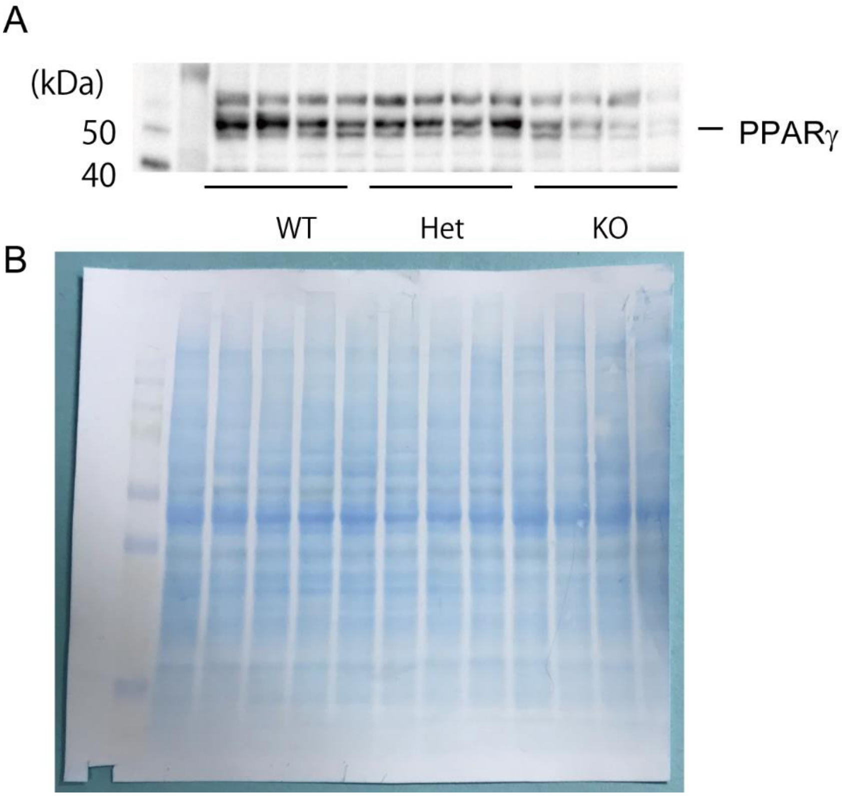


*A*, Western blotting image shown in Figure 2 (placental PPARγ abundance).

B, Coomassie brilliant blue-stained membrane used in *A*.

### Supplementary Tables

*Supplementary Table 1.*

Genotyping of adult mice from exon C-deletion heterozygous parents (Backcrossed onto B6 ; N=5)

| Sex | No. of mice | Genotype | | | *p*-value |
|---|---|---|---|---|---|
| | | +/+ | +/- | -/- | |
| Male | 82 | 32 | 39 | 11 | 0.004 |
| Female | 91 | 25 | 47 | 19 | 0.64 |
| Total | 173 | 57 | 86 | 30 | 0.015 |

*p* = Hardy-Weinberg's law test

*Supplementary Table 2.*

Genotyping of adult mice from exon C-deletion heterozygous parents (before backcrossing)

| Sex | No. of mice | Genotype | | | *p*-value |
|---|---|---|---|---|---|
| | | +/+ | +/- | -/- | |
| Male | 35 | 19 | 16 | 0 | < 0.001 |
| Female | 51 | 23 | 29 | 5 | 0.003 |
| Total | 86 | 42 | 45 | 5 | < 0.001 |

*p* = Hardy–Weinberg's law test

**Table S3. Materials used**

| Category | Material | Provider | Identifier |
|---|---|---|---|
| **Mice and transgenic lines** | | | |
| Mouse | C57BL/6J | Clea (Tokyo, Japan) | N/A |
| Vector | FRT-PGK-gb2-neo-FRTloxP | Gene Bridges GmbH, Heidelberg, Germany | A004 |
| Mouse | B6;CBA-Tg(CAG-Cre)47Imeg | Center for Animal Resources and Development, Kumamoto Univ., Kumamoto, Jap | CARD ID.272 |
| Mouse | B6;D2-Tg(CAG-Flp)18Imeg | Center for Animal Resources and Development, Kumamoto Univ., Kumamoto, Jap | CARD ID.265 |
| Instrument | Stereomicroscope | Nikon | SMZ745T with DS-L3 camera |
| **Quantitative RT-PCR** | | | |
| Reagent | SV Total RNA Isolation System | Promega, Japan (https://www.promega.jp/) | cat.no. Z3100 |
| Reagent | SuperScript IV VILO Master Mix | ThermoFisher Scientific(https://www.thermofisher.com/jp/ja/home.html) | cat.no. 11756050 |
| Reagent | THUNDERBIRD® SYBR qPCR mix | TOYOBO, http://lifescience.toyobo.co.jp/ | QPS-201 |
| Instrument | Real-Time PCR System | ThermoFisher Scientific(https://www.thermofisher.com/jp/ja/home.html) | QuantStudio™12K Flex |
| **Western blotting** | | | |
| Material | Polyvinylidene difluoride membranes (PVDF)/Immunobilon®-P | GE Healthcare | IPVH00010 |
| Antibody | Anti-PPARγ polyclonal antibody (1:1,000) | Santa Cruz | SC-7196 (H-100) |
| Antibody | Horseradish peroxidase-conjugated anti-rabbit IgG antibody (1:1,000) | GE Healthcare | NA934 |
| Antibody | Horseradish peroxidase-conjugated anti-mouse IgG antibody (1:1,000) | Sigma-Aldrich | A9044 |
| Reagent | ECL Prime™, Western Blotting Detection Reagents | GE Healthcare | RPN2232 |
| Reagent | MagicMark™ XP Western Protein Standard | ThermoFisher Scientific | LC5602 |
| Instrument | Luminescence imager | Bio-Rad laboratories | ChemiDoc™ MP system |
| **Histology** | | | |
| Instrument | Light microscopy | OLYMPUS Corp | BX53 with DP27 digital camera |
| Instrument | All-in-one fluorescent microscope | Keyence | BZ-X700 |
| **Immunohistochemistry** | | | |
| Reagent | Vector® M.O.M.™ Immunodetection Peroxidase Kit | Vector laboratories Inc. | PK-2200 |
| Antibody | Anti-PPARγ monoclonal antibody from mouse (1:500) | Perseus Proteomics, Inc. | A3409A |
| Antibody | Anti-monocarboxylate transporter 1 IgY antibody from Chiken (1:1,000) | Millipore | AB1286-I |
| Antibody | Alexa Fluor 488-conjugated goat anti-Chicken IgY (1:1,000) | ThermoFsher Scientific | A-11039 |
| **Software** | | | |
| | Photoshop CS5 | Adobe Systems | N/A |
| | ImageJ | https://imagej.nih.gov/ij/ | p1.52 |
| | JMP® | SAS Institute Inc. | ver. 13.2.1 |
| | PRISM® | GraphPad Software Inc. | ver 6.07 |

*Table S4. Primers used for quantitative RT-PCR*

| HGNC symbol | Accession No. | Forward primer (5'-3') | Reverse primer (5'-3') |
|---|---|---|---|
| **PPARγ** | | | |
| *Pparγ1* | NM_001127330.2 | CAGGACTGTGTGACAGACAAGAT | GGCCAGAATGGCATCTCTGTGTCAA |
| *Pparγ1sv* | AB644275 | GCGCTAAATTCTTCTTAACTC | GGCCAGAATGGCATCTCTGTGTCAA |
| *Pparγ2* | NM_011146 | GTTATGGGTGAAACTCTGGGAGAT | GGCCAGAATGGCATCTCTGTGTCAA |
| **Cyclins** | | | |
| *Ccna1* | NM_001305221.1 | ATGAGTTTGTCTACATCACCGACGA | TGATGCACACTCCTTGACGCCTT |
| *Ccna2* | NM_009828.3 | CGGAGCAAGAAAACCACTGACACC | GCTGCCTCTTCATGTAACCCACT |
| *Ccnb1* | NM_172301.3 | ATCCTCATTGACTGGCTAATACAGG | TGCAATAAACATGGCCGTTACACC |
| *Ccnb2* | NM_007630.2 | CTTACACCAGTTCCCAAATCCGAGA | GTCAGCTCCATCAGGTACTTGGCTA |
| *Ccnb3* | NM_183015.3 | AGTTCCTTCAGAATCCATTGCCACC | CTTGTCATCATTTGAAGCCACCGAT |
| *Ccnd1* | NM_007631.2 | AGGCGGATGAGAACAAGCAGA | CAGGCTTGACTCCAGAAGGG |
| *Ccnd2* | NM_009829.3 | GCGTGTTCGTCATCTGCTAGCC | CACCACATGCGTTACAACTATACGG |
| *Ccnd3* | NM_007632.2 | AGTTGCCAAAACGCCCCAGTACCTT | AATGACCACGGCACCCTTAAGACCC |
| *Ccne1* | NM_007633.2 | ACTTGGCACAGGACTTCTTTGATCGTT | ACATTCAGCCAGGACACAATGGTCA |
| *Ccne2* | NM_001037134.2 | ACAAAAGGAAAACAGATACGTGCAT | GCACCATCAGTGACGTAAGCAA |
| **E2F** | | | |
| *E2f1* | NM_007891.5 | GGGCTGGGTTTGAAACTCTC | GAGTGAACATTCCCTCCAACA |
| *E2f2* | NM_177733.7 | TCGCTTTACACGCAGACG | GCACATCGCACAATTTGG |
| *E2f6* | NM_033270.2 | TTCGGAAGAGGCGAGTGTAT | CTTTTCCACCAGTTCGATGC |
| *E2f7* | NM_178609.4 | CTAAAGTCTGGGTGCCTTGTG | CAGTGTGACCTCATAGTTCATCG |
| *E2f8* | NM_001013368.5 | GGCCACCAACCATGACTC | GCGACTGGTTGTCCGTTTA |
| **CDK inhibitors** | | | |
| *p19* | NM_009878 | AGCCTTACTGGGTTACTTGTCAACA | CTGTAGGAGCCCCTTCTTTGTCCA |
| *p21* | NM_007669.5 | CTGGTTCCTTGCCACTTCTTACCTG | TTACGGTTGAGTCCTAACTGCCAT |
| *p27* | NM_009875.4 | GTCGCAGAACTTCGAAGAGG | AAACCGAACAAAAGCGAAAC |
| *p57* | NM_001161624.1 | CTGAAGGACCAGCCTCTCTC | TGCTCTACGCAACCATCTCC |
| **Pregnancy-specific glycoproteins** | | | |
| *Psg16* | NM_007676.4 | CTCCAATAGTGACACCTAACCCCAA | AAACTGTGAATCAAAACTATCTAGTAGCCA |
| *Psg19* | NM_011964.2 | TCCAGTGCCACCACATGCTGTC | TGCACGGCCACTGATGATAGACTCT |
| *Psg21* | NM_027403.4 | TTCTCCCACATCCCCTCTC | GGGGAAAATAATAAGTGGAAGCA |
| *Psg22* | NM_001004152.2 | CACAGTGGAAGAGAGATATTGTTCA | AAGCCAGAGTCTTTCTCAGTGAC |
| *Psg23* | NM_020261.4 | GAGCCTGTCCCCGTCAAAGTGT | GAAATGCCTCTGCCCTGCTATAGT |
| **Housekeeping genes** | | | |
| *18s* | NR_003278.3 | CGGCTACCACATCCAAGGAA | GCTGGAATTACCGCGGCT |
| *HPRT* | NM_013556.2 | CTATAAGTTCTTTGCTGACCTGCT | ATCATCTCCACCAATAACTTTTATGT |
